# Evolution of metabolic novelty: a trichome-expressed invertase creates specialized metabolic diversity in wild tomato

**DOI:** 10.1101/502971

**Authors:** Bryan J. Leong, Daniel Lybrand, Yann-Ru Lou, Pengxiang Fan, Anthony L. Schilmiller, Robert L. Last

## Abstract

Plants produce myriad taxonomically restricted specialized metabolites. This diversity – and our ability to correlate genotype with phenotype – makes the evolution of these ecologically and medicinally important compounds interesting and experimentally tractable. Trichomes of tomato and other nightshade family plants produce structurally diverse protective compounds termed acylsugars. While cultivated tomato (*Solanum lycopersicum*) accumulates strictly acylsucroses, the South American wild relative *Solanum pennellii* produces copious amounts of acylglucoses. Genetic, transgenic and biochemical dissection of the *S. pennellii* acylglucose biosynthetic pathway identified a trichome gland cell expressed invertase-like enzyme that hydrolyzes acylsucroses (Sopen03g040490). This enzyme acts on the pyranose ring-acylated acylsucroses found in the wild tomato but not the furanose ring-decorated acylsucroses of cultivated tomato. These results show that modification of the core acylsucrose biosynthetic pathway leading to loss of furanose ring acylation set the stage for co-option of a general metabolic enzyme to produce a new class of protective compounds.

## Introduction

Plants synthesize hundreds of thousands of structurally diverse and lineage-, tissue-, or cell type-specific specialized metabolites (*1*). These compounds act as anti-herbivory, antimicrobial, or allelopathic compounds (*2–4*). The enzymes that produce these specialized metabolites arise from a variety of proteins through gene duplication and neo- or sub-functionalization. Examples include diversification of existing specialized metabolic enzymes (*5*) and co-option of core metabolic activities (*6*). In the latter mechanism, gene duplication can remove stabilizing selective pressure on one of the paralogs associated with core metabolism, enabling evolution of the novel enzymatic activity without disrupting growth and development. This leads to new and modified pathways that produce structurally and functionally diverse specialized metabolites.

Glandular trichome-synthesized acylated sugars (‘acylsugars’) are structurally diverse specialized metabolites found throughout the Solanaceae (*7–13*). These compounds have documented roles in direct and indirect protection against herbivores and microbes (*14*, *15*), as well as allelopathic properties (*15, 16*). Their low toxicity to vertebrates generates interest in generating plant breeding strategies for deploying acylsugars in crop protection (*17*, *18*). These metabolites consist of a sugar core – typically sucrose – with aliphatic chains of variable length, structure and number attached by ester linkages. Acylsugars were reported from genera across the Solanaceae family, including *Datura, Nicotiana, Petunia, Physalis, Salpiglossis*, and *Solanum* with single species producing at least three dozen chromatographically distinct acylsugars (*10*, *13, 16, 19–22*).

In recent years, several evolutionarily-related enzymes were implicated in the core acylsucrose biosynthetic pathways in species across the family, including the cultivated tomato *Solanum lycopersicum, Petunia axillaris* and *Salpiglossis sinuata* (*7*, *9–12, 23*). These biosynthetic pathways consist of trichome-expressed BAHD-family acylsugar acyltransferases (ASATs) (*7, 9, 23*), which sequentially transfer acyl groups from acyl-coenzyme A (acyl-CoA) substrates to specific hydroxyl groups of sucrose (*7, 9, 23*).

The cultivated tomato biosynthetic network is well characterized, with four ASATs – S1ASAT1 through SlASAT4 – catalyzing consecutive reactions to produce tri- and tetra-acylated sucroses. S1ASAT1 acts first by transferring an acyl chain to the R4 hydroxyl of the pyranose ring of sucrose, and SlASAT2 transfers an acyl chain to the R3 position of the monoacylated sucrose (*23*). Next, SlASAT3 acylates the diacylated sucroses at the furanose ring R3’ position (*9*). SlASAT4 completes the pathway by transferring an acetyl group to the pyranose ring R2 position of a triacylsucrose (*7, 22*). Enzyme promiscuity and the presence of an array of acyl-CoAs results in production of a diverse group of acylsucroses in *S. lycopersicum* (*11*, *24*).

The metabolic diversity in acylsugars is even greater in the broader *Solanum* genus. The wild relative of tomato, *S. pennellii* LA0716, is a prime example, producing a mixture of abundant acylsucroses that are distinct from those in *S. lycopersicum*. While *S. lycopersicum* accumulates acylsucroses with two or three acylations on the pyranose ring and a single acylation at the furanose ring R3’ position (termed ‘F-type’ acylsucroses), *S. pennellii* accumulates distinct ‘P-type’ triacylsucroses acylated only on the pyranose R2, R3, and R4 positions. P-type acylsucroses are synthesized by *S. pennellii* orthologs of the *S. lycopersicum* ASAT1, ASAT2, and ASAT3 enzymes (Fig. 1A). The different acylation pattern observed in *S. pennellii* results from altered substrate specificity and acylation position of SpASAT2 and SpASAT3 relative to their *S. lycopersicum* counterparts (*11*).

**Fig. 1.**
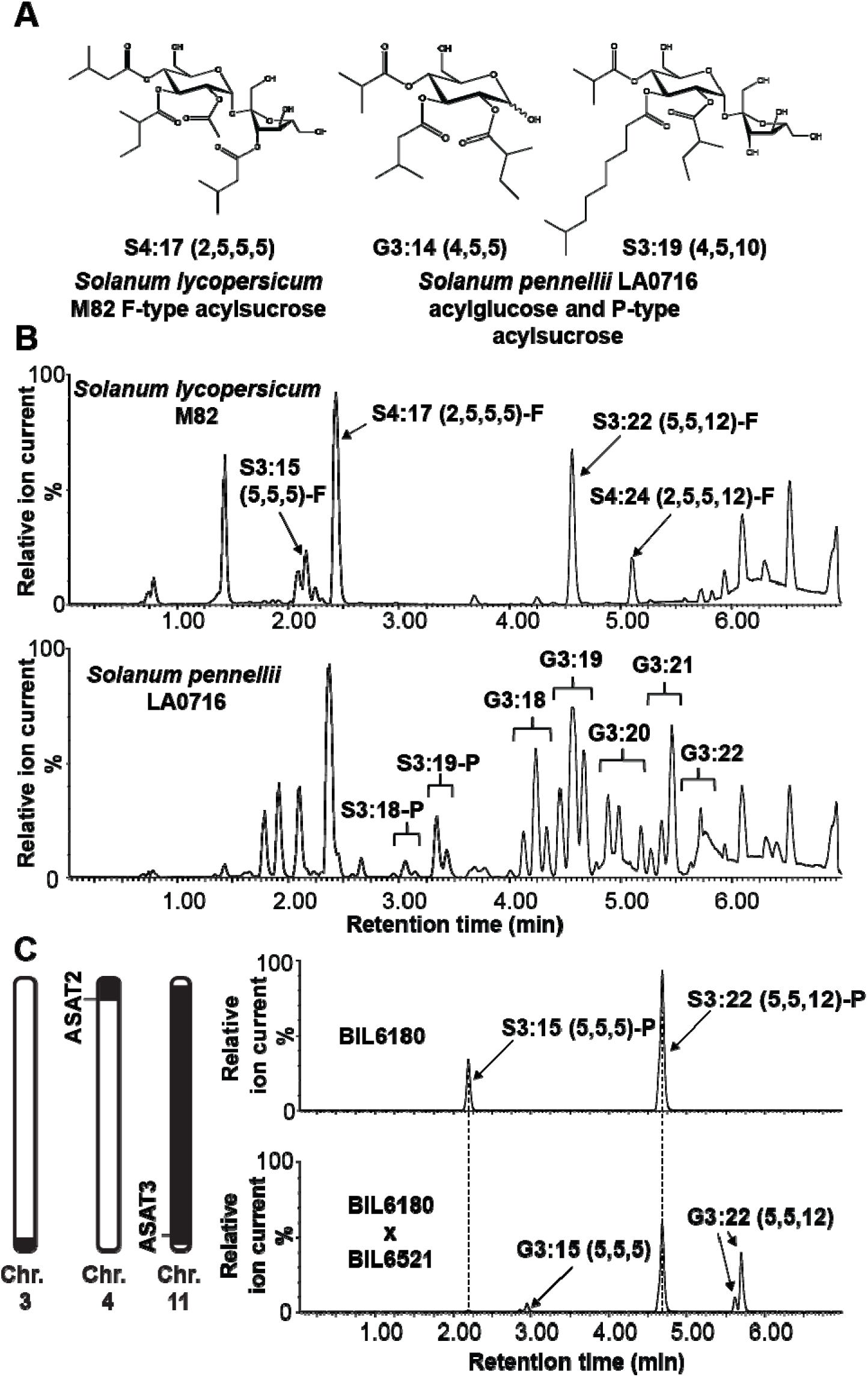
Three *S. pennellii* LA0716 regions condition acylglucose accumulation. (**A**) Examples of NMR-resolved *S. lycopersicum* and *S. pennellii* acylsugar structures. Acylsugars from *S. lycopersicum* are composed of sucrose acylated on both the pyranose and furanose rings (‘F-type’). *S. pennellii* acylsugars are a mixture of sucrose (‘P-type’) and glucose-based compounds with acylation exclusively on the pyranose ring. (**B**) Acylsugar ESI-mode LC-QToF MS profiles. Top: *S. lycopersicum* M82 with acylsucroses S3:15 (5,5,5)-F, S4:17 (2,5,5,5)-F, S3:22 (5,5,12)-F, and S4:24 (2,5,5,12)-F annotated. Bottom: S. *pennellii* LA0716 acylsucroses and acylglucoses. (**C**) Left: Representation of *S. pennellii* chromosomal introgressions in BIL6521 x BIL6180 progeny that contain QTLs affecting acylglucose biosynthesis (*30*). The black portions of the chromosomes correspond to *S. pennellii* introgressions, while the white portions correspond to the chromosomal regions in the M82 background. Right: ESI-mode LC-MS analysis of BIL6180 compared with the BIL6180 x BIL6521 F2 progeny reveals acylglucose accumulation in the hybrid, but not in BIL6180.

*S. pennellii* LA0716 has other acylsugar characteristics that differentiate it from cultivated tomato (Fig. 1B). First, it produces copious amounts of acylsugars that render the plant sticky, representing up to ~20% of leaf dry weight (*19, 25*). Second, the vast majority of *S. pennellii* LA0716 acylsugars are glucose molecules with three acyl chains (termed “acylglucoses”) (Fig. 1A and B), while only 7–16% of total acylsugars are acylsucroses (*26*). In contrast to the well-characterized *S. pennellii* acylsucrose biosynthetic enzymes (*9, 11*), no complete acylglucose metabolic pathway has yet been described. This is despite the fact that acylglucoses were also characterized in several additional Solanaceae species (*16*, *20*). A previously proposed partial *S. pennellii* pathway invoked two glucosyltransferases capable of creating 1-*O*-acyl-D-glucose from UDP-glucose and free fatty acids of differing structures (*27*). This mechanism proposed a second step in which a serine carboxypeptidase-like (SCPL) acyltransferase catalyzed disproportionation of two 1-*O*-isobutyrl-D-glucose molecules to yield one 1,2-*O*-di-isobutyrl-D-glucose (*28, 29*). However this pathway is unlikely to function *in vivo* as the 1,2,-*O*-diacylglucoses obtained *in vitro* differ from the 2,3,4-*O*-tri-acylglucoses observed in *S. pennellii* both in number (two instead of three) and position of acyl chains: *S. pennellii* acylglucoses bear chains at the R2, R3, and R4 positions rather than at position R1 (*19*).

In contrast to the unsubstantiated published biosynthetic pathway, compelling quantitative trait locus (QTL) and biochemical results implicate multiple genetic loci in acylglucose accumulation in *S. pennellii* LA0716. The combination of three *S. pennellii* regions on chromosomes 3, 4, and 11 cause *S. lycopersicum* breeding line CU071026 to accumulate acylsugars comprising up to 89% acylglucoses (*30*). The presence of QTLs on both chromosomes 3 and 11 yields detectable acylglucoses, while addition of the chromosome 4 locus leads to elevated accumulation. Notably, the chromosome 4 and 11 QTLs respectively include the *SpASAT2* and *SpASAT3* genes responsible for accumulation of P-type acylsucroses in *S. pennellii* (*9, 23*). The agreement of QTL and biochemical data is consistent with the hypothesis that SpASAT2 and SpASAT3 produce P-type acylsucroses that are substrates for a chromosome 3 factor that then synthesizes triacylglucoses.

We report the characterization of the plant specialized metabolic invertase-like enzyme **a**cyl**s**ucrose **f**ructo**f**uranosidase **1** (SpASFF1; Sopen03g040490), a chromosome 3 β-fructofuranosidase capable of cleaving the glycosidic bond of P-type acylsucroses. Genetic and transgenic plant approaches demonstrate that *S. pennellii* LA0716 acylglucose production requires *SpASFF1*. This work also documents a three-gene epistatic interaction between the *SpASAT2, SpASAT3*, and *SpASFF1* loci that conditions high-level acylglucose accumulation. While yeast invertase and other variants involved in core metabolism have been studied since the 19^th^ century (*31*–*33*), this work documents a new type of role for β-fructofuranosidase type enzymes in specialized metabolism. These results extend our understanding of evolutionary mechanisms leading to trichome specialized metabolic diversity by demonstrating how neofunctionalization led to co-option of invertase from general metabolism into a cell-type-specific specialized metabolic network.

## Results

### A *S. pennellii* chromosome 3 locus is necessary for acylglucose production from P-type acylsucroses

Published QTL mapping studies indicate that introgression of *S. pennellii* LA0716 loci on chromosomes 3, 4 and 11 leads to accumulation of acylglucoses in a cultivated tomato *S. lycopersicum* background (*30*, *34*). Three introgression lines harboring individual acylglucose QTLs in the *S. lycopersicum* background were screened, but none of the single introgressions in lines IL3-5, IL4-1, or IL11-3 (*35*) yielded detectable leaf acylglucoses (Fig. S1). These observations are consistent with the hypothesis that multiple *S. pennellii* loci are needed for *S. lycopersicum* acylglucose accumulation. Indeed, there are low but detectable levels of acylglucoses (87% of total acylsugars; Fig. S2) in backcross inbred line BIL6521 (*36*), which contains *S. pennellii* LA0716 introgressions from chromosomes 1, 3, and 11. This BIL accumulates four acylglucoses (Table S1), with the major one, G3:22 (5,5,12) (Fig. S3 and Fig. S4), resembling the pyranose ring of the P-type acylsucrose S3:22 (5, 5, 12)-P detected at low levels in trichomes of the single chromosome 11 introgression line, IL11-3 (*9*). In fact, BIL6521 accumulates a P-type acylsucrose, S3:22 (Fig. S3 and Fig. S4). These results are consistent with the hypothesis that the chromosome 3 region is necessary for acylglucose production, but only when P-type acylsucroses are produced. Note that in our nomenclature, ‘S’ and ‘G’ refer to sucrose or glucose core, respectively, and 3:22 (5,5,12) indicates that there are three ester-linked acyl chains of 5, 5 and 12 carbons, for a total of 22 chain carbons (*9*). When NMR-derived structural information is available, superscripts indicate acyl chain positions with R representing the pyranose ring, and R’ representing the furanose ring (Fig. 1A).

Because the acylsugar levels in BIL6521, which lack a chromosome 4 introgression, were much lower than most other lines, we tested the impact of adding a chromosome 4 introgression carrying the *SpASAT2* locus. A cross was made between BIL6521 and BIL6180, a recombinant line harboring introgressions on chromosomes 4, 5 and 11, which includes both the *SpASAT2* and *SpASAT3* loci (Fig. 1C). BIL6180 was previously found to produce only P-type acylsucroses as a result of the chromosome 4 and 11 introgressions (Fig. 1C), however, it accumulated significantly higher overall levels of acylsucroses compared to BIL6521 (Fig. S2) as well as other short-chain containing P-type acylsucroses not present in BIL6521. If all P-type acylsucroses are substrates for a *S. pennellii* LA0716 factor on chromosome 3, we predicted that both of the corresponding acylglucoses G3:15 (5,5,5) and G3:22 (5,5,12) would accumulate in a line harboring the chromosome 3, 4 and 11 introgressions. Indeed, the F2 progeny of BIL6521 x BIL6180, genotyped as heterozygous for the *S. pennellii* chromosome 3, 4 introgressions, and homozygous for the *S. pennellii* chromosome 11 region, produced these two predicted acylglucoses (Fig. 1C, Fig. S3 and Fig. S4). These findings – in combination with the published QTL results – indicate that the *S. pennellii* chromosome 3 introgression is necessary for acylglucose biosynthesis and suggests that P-type acylsucroses are acylglucose biosynthetic precursors.

### The chromosome 3 locus encodes a glandular trichome-expressed β-fructofuranosidase

We sought candidate glycoside hydrolase genes in the 1.7-Mb QTL AG3.2, the acylglucose-associated region from *S. pennellii* LA0716 previously mapped to the bottom of chromosome 3 (*30*) (Fig. 2A). Three of the 238 genes in this region of the *S. lycopersicum* Heinz 1706 genome assembly SL2.50 annotation (*37*) are predicted as encoding glycoside hydrolases (members of the GH32, GH35, and GH47 families; Table S2). We focused on the GH32 family Sopen03g040490 gene because all previously characterized members of the family have β-fructofuranosidase or fructosyltransferase activity (*38*). As acylsucroses are β-fructofuranosides, we hypothesized that the GH32 enzyme cleaves the glycosidic bond of P-type acylsucroses to generate acylglucoses. Based on the full results of this study, we designate this gene *<u>ACYLS</u>UCROSE <u>F</u>RUCTO<u>F</u>URANOSIDASE1* (*ASFF1*).

**Fig. 2.**
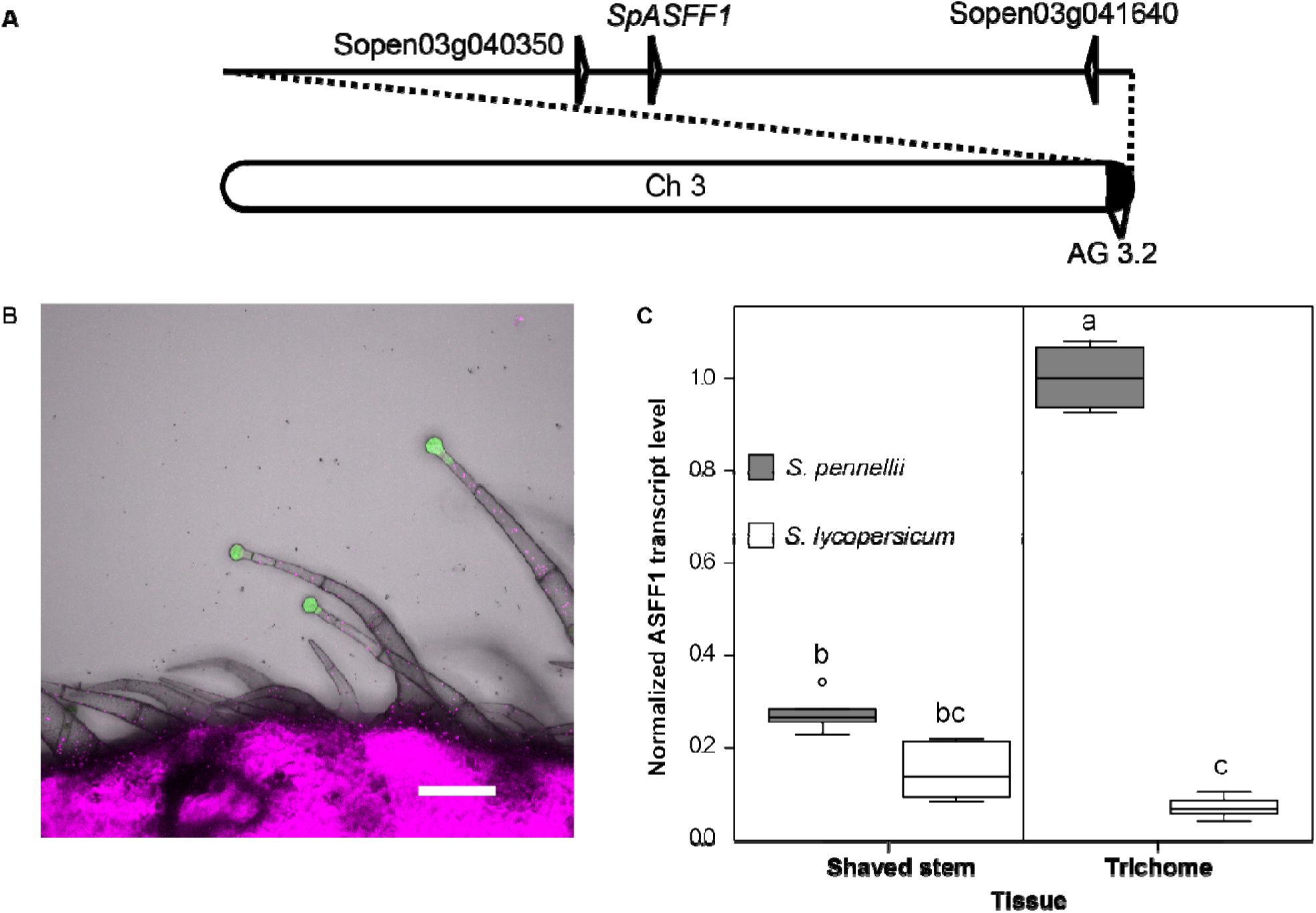
Glycoside hydrolase 32 family gene *SpASFF1* from QTL AG 3.2 shows trichome-specific expression. (**A**) Chromosome 3 with the AG3.2 introgression (*30*). Positions of three glycoside hydrolase genes (*Sopen03g040350, Sopen03g041640*, and *SpASFF1*) are indicated. (**B**) Expressing GFP-GUS under control of the native *ASFF1* promoter from *S. pennellii* LA0716 yields GFP signal in *S. lycopersicum* M82 type IV trichome tip cells, but not stalk cells or stem tissue. Green channel indicates GFP signal; magenta channel shows chlorophyll fluorescence. Scale bar = 100 μm. (**C**) qRT-PCR analysis of ASFF1 transcripts shows statistically significantly higher levels in *S. pennellii* LA0716 trichomes compared to underlying stem tissue or trichomes of *S. lycopersicum* M82. Treatments that do not share a letter are significantly different from one another (*p* < 0.001; one-way ANOVA, Tukey’s Honestly Significant Difference mean-separation test). Whiskers represent minimum and maximum values less than 1.5 times the interquartile range from the 1^st^ and 3^rd^ quartiles, respectively. Values outside this range are represented as circles; *n* = 6 for all species and tissue types.

*S. lycopersicum* acylsugars accumulate in type I/IV glandular trichome tip cells (*39*) and trichome tip cell-specific gene expression is a hallmark of all characterized acylsugar biosynthetic genes (*e.g., ASAT1/2/3/4, IPMS3*) (*7, 9, 23, 24*). We used a reporter gene approach to ask whether *SpASFF1* exhibits trichome-specific expression. The 1.8-kb region immediately upstream of the *SpASFF1* ORF in the *S. pennellii* LA0716 genome drove expression of a green fluorescent protein-β-glucuronidase fusion protein (GFP-GUS) in *S. lycopersicum* M82 plants. Indeed, GFP signal in transformed plants was observed in the tip cells of type I/IV trichomes but not in the trichome stalk cells or underlying stem epidermis (Fig. 2B). This result is consistent with a role of SpASFF1 enzyme in type I/IV trichome metabolism.

We cross-validated the trichome enriched expression pattern of *ASFF1* in *S. pennellii* LA0716 using Quantitative Reverse Transcriptase PCR (qRT-PCR). ASFF1 transcript levels were 3.7-fold higher in trichomes of *S. pennellii* LA0716 stems than in underlying shaved stem tissue (*p* < 0.001; one-way ANOVA, Tukey’s Honestly Significant Difference (HSD) mean-separation test) (Fig. 2C). The observed enrichment of transcripts in trichome samples is similar to analysis of previously identified acylsugar biosynthetic genes from tomato, petunia and tobacco (*7, 12, 15, 23, 24*). Together, transcript enrichment in trichomes and restriction of gene expression to trichome tip cells support the hypothesis that *SpASFF1* acts in acylsugar biosynthesis.

Acylglucoses accumulate in *S. pennellii* LA0716 but not in *S. lycopersicum* M82 (*8, 19*). However, *ASFF1* is predicted to encode a full open reading frame in both the *S. lycopersicum* Heinz 1706 and the *S. pennellii* LA0716 genomes (*37, 40*). While *ASFF1* transcripts are enriched in trichomes of *S. pennellii* LA0716, no significant difference was observed between *ASFF1* transcript levels in *S. lycopersicum* M82 stems and trichomes (*p* = 0.063; one-way ANOVA, Tukey’s HSD mean-separation test) (Fig. 2C). Additionally, we found that *ASFF1* transcripts are enriched 14-fold in *S. pennellii* LA0716 trichomes relative to transcripts in trichomes of *S. lycopersicum* M82 (*p* < 0.001; one-way ANOVA, Tukey’s HSD mean-separation test) (Fig. 2C). Together, the tissue- and species-level specificity of *ASFF1* expression is consistent with a role for the gene in acylglucose biosynthesis.

### Gene editing reveals that *SpASFF1* is necessary for *S. pennellii* LA0716 acylglucose accumulation

We used CRISPR/Cas9-mediated gene editing in *S. pennellii* LA0716 to test whether *SpASFF1* is necessary for acylglucose accumulation. Two small guide RNAs (sgRNAs) targeting the third *SpASFF1* exon were used to promote site specific DNA cleavage by hCas9 in the stably transformed plants (Fig. 3A and Fig. S5). Three homozygous T_1_ mutants were obtained with different site-specific mutations, each of which is predicted to cause complete loss of function through translational frame-shifts and premature protein termination. Two of them (*spasff1-1-1* and *spasff1-1-2*), which carry 228 bp and 276 bp insertion-deletions, respectively, are derived from segregation of one heteroallelic T_0_ plant. The third mutant (*spasff1-2*) with a 1 bp insertion is the descendant of a homozygous T_0_ mutant.

**Fig. 3.**
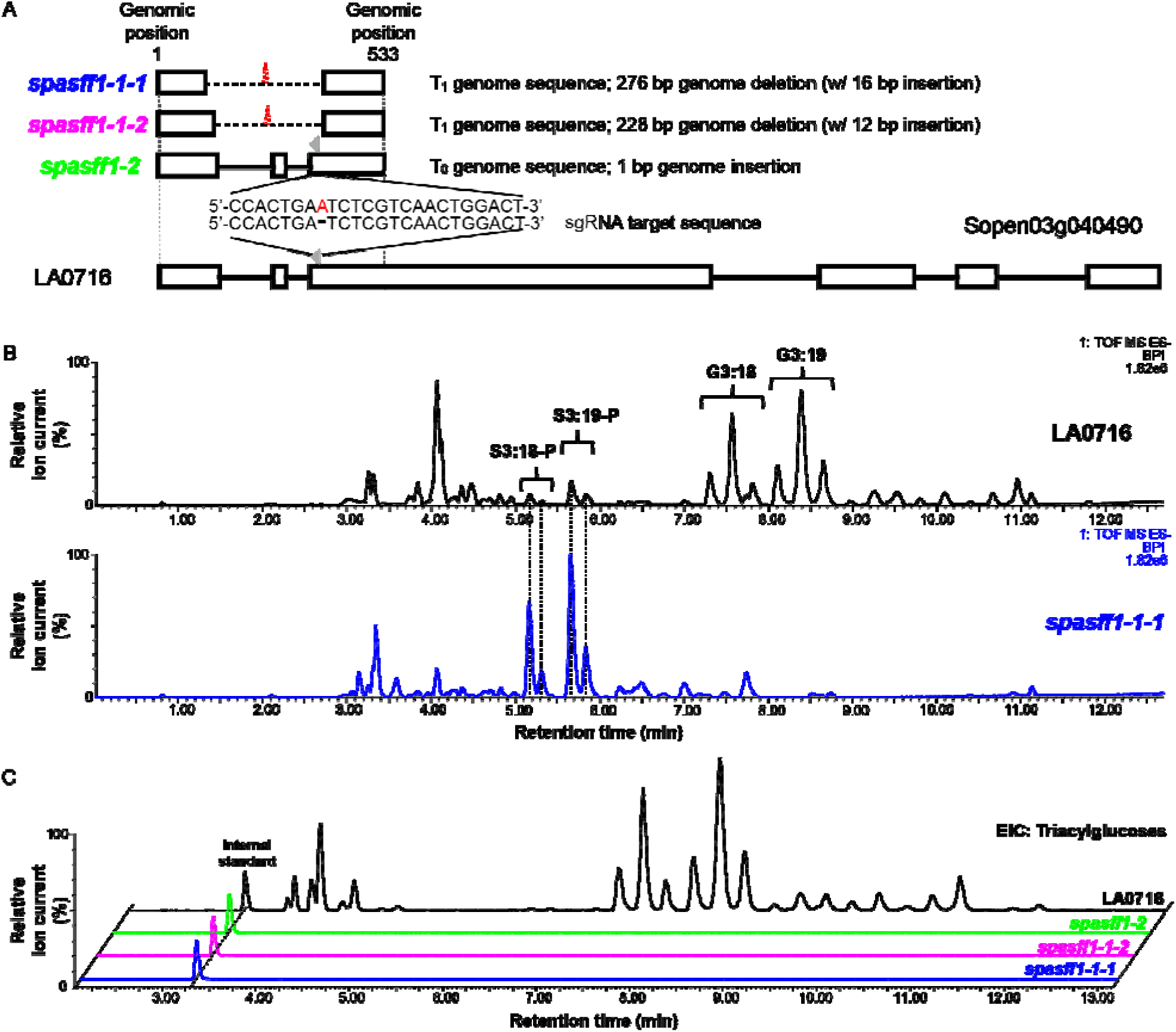
CRISPR/Cas9-mediated *S. pennellii* LA0716 *SpASFF1* knockouts eliminate detectable acylglucoses. (**A**) Schematic representation of mutagenesis strategy with two sgRNAs (grey arrowheads) targeting the *Spasff1* ORF that result in three homozygous knockout lines. White boxes indicate exons, horizontal bars indicate introns, dotted lines indicate deletions and red letter indicates insertion. Mutant allele DNA sequences are found in Fig. S5. (**B**) Mutant line *spasff1-1-1* accumulates abundant acylsucroses but no detectable acylglucoses. ESI-base-peak intensity LC-MS chromatograms are shown for *spasff1-1-1* and LA0716. (**C**) Triacylglucose extracted ion chromatograms of trichome extracts from *S. pennellii* LA0716 and three *spasff1* mutant plants show that homozygous *asff1* lines produce undetectable levels of triacylglucose. Extracted ion chromatogram values displayed: G3:12 (*m/z:* 435.19), G3:13 (*m/z:* 449.2), G3:14 (*m/z:* 463.22), G3:15 (*m/z:* 477.23), G3:16 (*m/z:* 491.28), G3:17 (*m/z:* 505.26), G3:18 (*m/z:* 519.28), G3:19 (*m/z:* 533.30), G3:20 (*m/z:* 547.31), G3:21 (*m/z:* 561.33), G3:22 (m/z: 575.34), and telmisartan (internal standard) (*m/z*:513.23). ESI-mode Base peak intensity (BPI) LC-MS chromatograms for all lines are shown in Fig. S6. Note: For panel B and C, *spasff1-1-1/1-1-2* are homozygous T2 lines, while *spasff1-2* are homozygous T1 lines that were all grown together. *spasff1* lines were diluted 100 fold before LC-MS analysis to avoid saturation of the LC-MS detector. This is due to differences in ionization between acylsucroses and acylglucoses in ESI-mode.

Results from LC-MS analysis of leaf surface metabolites from these lines were consistent with the hypothesis that *SpASFF1* is necessary for acylglucose biosynthesis. All *spasff1* lines failed to accumulate detectable acylglucoses (Fig. 3B and C and Fig. S6), but produced acylsucroses at levels comparable to total acylsugars in wild-type *S. pennellii* plants (Table 1).

**Table 1.** Quantitative analysis of acylsugars from *S. pennellii* LA0716 and three *Spasff1* lines. Acyl chains were saponified from the acylsugars and the resulting sugar cores analyzed by UPLC-ESI-Multiple Reaction Monitoring. Data are shown from individual T1 homozygous plants grown together but independently from those in Fig. 3. Note: These extracts include other glycosylated compounds such as flavonoids, which could be responsible for the non-zero values for glucose measurements in plants lacking detectable acylglucoses (*61*).

| Line | Plant number | Sucrose (%) | Glucose (%) | Total sugar core (nmol/mg DW) |
| --- | --- | --- | --- | --- |
| LA0716 | 1 | 1% | 99% | 41.93 |
|  | 2 | 3% | 97% | 15.78 |
|  | 3 | 1% | 99% | 33.70 |
|  | 4 | 2% | 98% | 55.89 |
|  | 5 | 1% | 99% | 138.80 |
|  | 6 | 1% | 99% | 29.34 |
| *spasff1-1-1* | 1 | 98% | 2% | 25.82 |
|  | 2 | 99% | 1% | 23.23 |
|  | 3 | 98% | 2% | 26.79 |
|  | 4 | 99% | 1% | 21.49 |
| *spasff1-1-2* | 1 | 99% | 1% | 18.74 |
|  | 2 | 99% | 1% | 24.55 |
|  | 3 | 99% | 1% | 28.72 |
|  | 4 | 99% | 1% | 18.35 |
| *spasff1-2* | 1 | 97% | 3% | 80.01 |
|  | 2 | 98% | 2% | 37.51 |
|  | 3 | 95% | 5% | 84.44 |

### SpASFF1 converts pyranose ring-acylated P-type acylsucroses to acylglucoses both *in vivo* and *in vitro*

The results described above strongly suggest that SpASFF1 converts pyranose ring-acylated P-type acylsucroses to acylglucoses. In addition, IL3-5 does not accumulate detectable acylglucoses despite possessing *S. pennellii ASFF1*, suggesting that F-type acylsucroses are not substrates for SpASFF1 (Fig. 4A and Fig. S7). We took a transgenic approach to ask whether *SpASFF1* alone is sufficient to confer acylglucose accumulation in a P-type acylsucrose-accumulating background. We transformed the P-type acylsucrose-producing *S. lycopersicum* double introgression BIL6180 (Fig. 4B) with a T-DNA containing the open reading frame of *SpASFF1* and the 1.8 kb immediately upstream of its start codon (Fig. 4C). In addition to the P-type acylsucroses in the parental BIL6180, the *SpASFF1* transgenics accumulated major hexose acylsugars with MS characteristics consistent with G3:15 (5,5,5) and G3:22 (5,5,12) (Fig. 4C, Fig. S8). The acyl chain composition of these acylglucoses matches the S3:15 (5^R2^,5^R3^,5^R4^) and S3:22 (5^R2^,5^R4^,12^R3^) P-type acylsucroses detected in BIL6180 (Fig. S4). Acylglucoses in the transgenic lines are also identical to those seen in BIL6521 x BIL6180 based on LC retention time and MS fragmentation. This confirms that SpASFF1 converts *S. pennellii* P-type acylsucroses produced by SpASAT2 and SpASAT3 to acylglucoses. Taken together, these *in vivo* results indicate that the presence of *SpASFF1* is sufficient to yield acylglucoses *in vivo* when P-type acylsucroses are present, but not in plants accumulating only F-type acylsucroses.

**Fig. 4.**
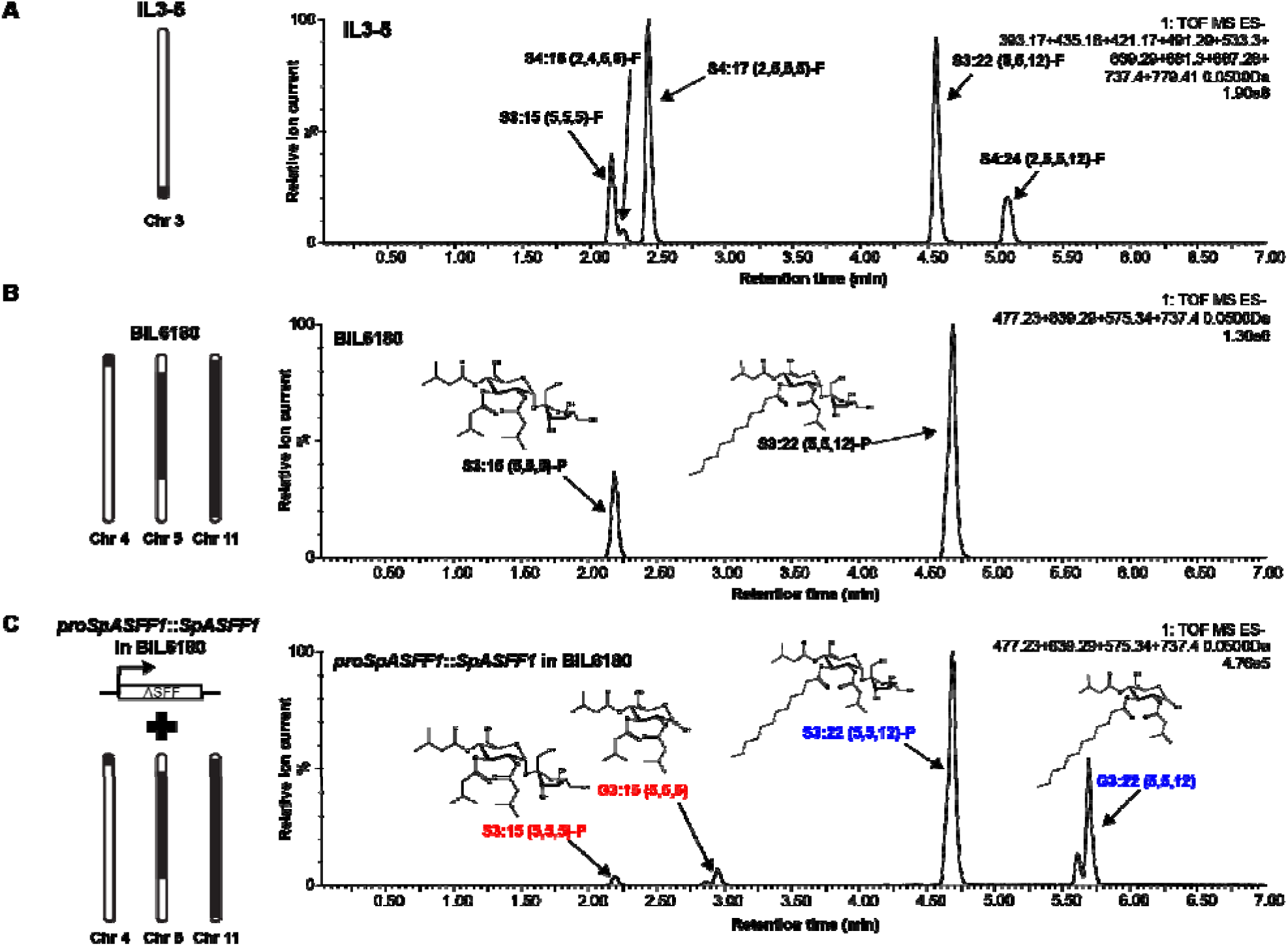
Expression of *SpASFF1* in P-type acylsucrose producing BIL6180 trichomes results in accumulation of acylglucoses in surface extracts. (**A**) IL3-5 accumulates F-type acylsucroses without detectable acylglucoses. ESI-mode LC-MS analysis of trichome extracts of IL3-5 are shown. Extracted ion chromatograms of S3:15 (*m/z:* 639.29), S4:16 (*m/z:* 667.28), S4:17 (*m/z:* 681.30), S3:22 (*m/z:* 737.40), and S4:24 (*m/z:* 779.41) in addition to their glucose cognates (missing a C5 chain present on the furanose ring), G2:10 (m/z: 393.17), G3:11 (*m/z:* 421.17), G3:12 (*m/z:* 435.18), G2:17 (*m/z:* 491.29), and G3:19 (*m/z:* 533.30) are shown. (B) BIL6180 accumulates P-type acylsucroses with no detectable acylglucoses. ESI-mode LC-MS analysis of trichome extracts of BIL6180 are shown. Extracted ion chromatograms of S3:15 (*m/z*: 639.29), S3:22 (*m/z*:737.40), G3:15 (*m/z*: 477.23), and G3:22 (*m/z:* 575.34) are shown. (**C**) Introduction of *SpASFF1* driven by its endogenous promoter in BIL6180 is sufficient to cause accumulation of detectable G3:15 and G3:22 acylglucoses. ESI-mode LC-MS analysis of trichome extracts of a *proSpASFF1::SpASFF1* in a BIL6180 T_2_ line is shown. Extracted ion chromatograms of S3:15 (*m/z:* 639.29), S3:22 (*m/z*:737.40), G3:15 (*m/z:* 477.23), and G3:22 (*m/z:* 575.34) are shown. Note: All *m/z* values correspond to the formate adducts of those acylsugars. Mass window: 0.05Da in all experiments. Acylglucose structure is inferred from collision induced dissociation-mediated fragmentation (Fig S8).

*In vitro* assays supported the hypothesis that SpASFF1 accepts P-type, but not F-type acylsucroses as substrates. Initial attempts to express SpASFF1 fusion proteins in *E. coli* did not produce soluble protein. For this reason, recombinant His-tagged SpASFF1 was expressed using the *Nicotiana benthamiana* transient expression system (*41*). The enzyme was tested with both P-type and F-type acylsucrose substrates purified from *S. pennellii asff1* and *S. lycopersicum* M82, respectively. Consistent with *in vivo* observations, SpASFF1 demonstrated hydrolytic activity with purified P-type S3: 19 (4^R4^,5^R2^,10^R3^) (*42*), yielding a compound with *m/z* consistent with a G3:19 (4,5,10) structure (Fig. 5A and Fig. S9). In contrast, SpASFF1 demonstrated no hydrolytic activity with F-type S3:22 (5^R4^,5^R3’^,12^R3^) (*8*) (Fig. 5B), suggesting that the presence of an acyl chain on the sucrose furanose ring prevents enzymatic hydrolysis. We further observed that SpASFF1 activity was undetectable with unmodified sucrose, while a commercially available yeast invertase hydrolyzed sucrose, but not S3:19 (Fig. S10). This SpASFF1 *in vitro* substrate specificity corroborates the *in vivo* results showing that acylglucoses only accumulate in lines containing P-type acylsucroses.

**Fig. 5.**
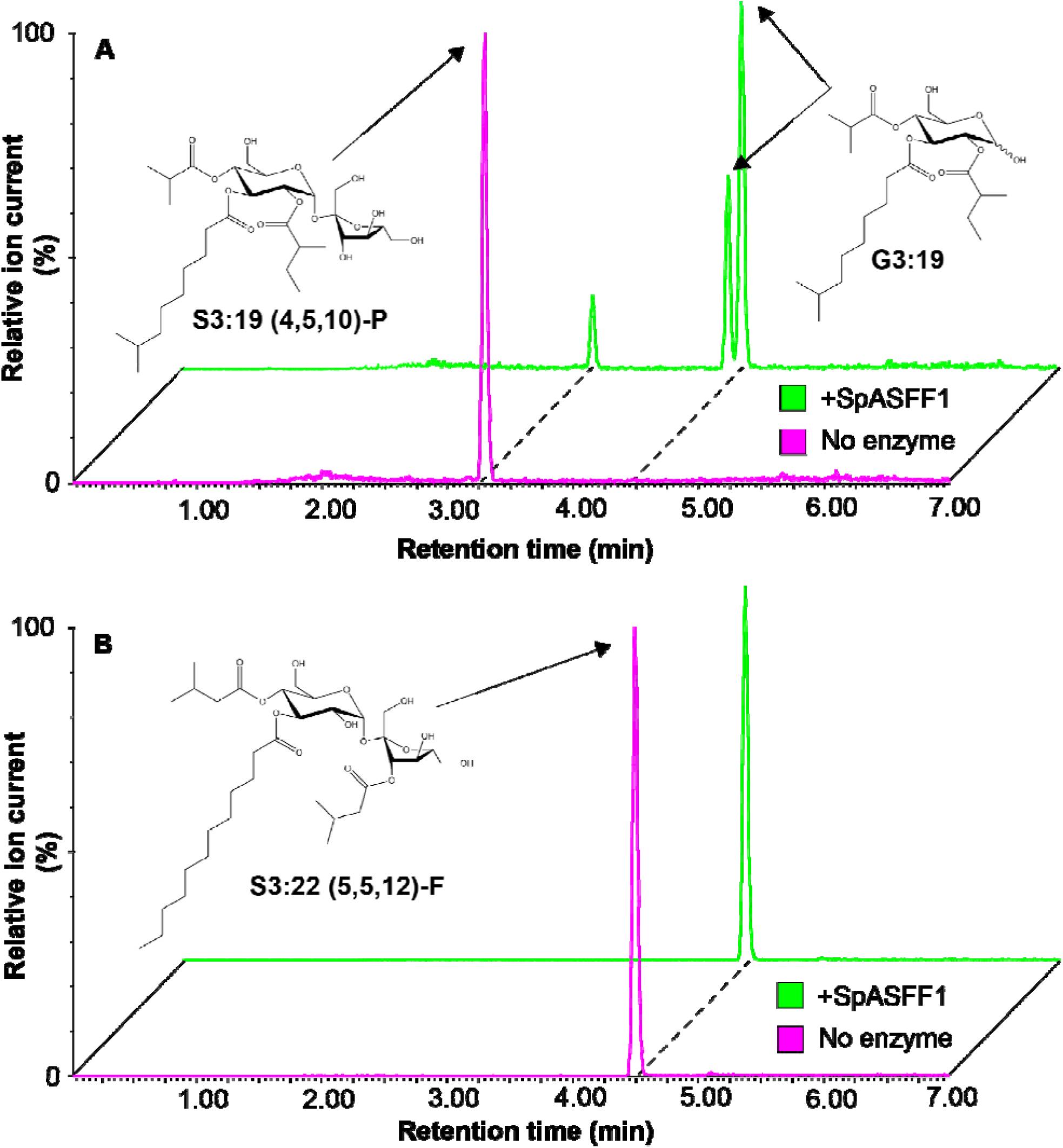
SpASFF1 cleaves a P-type S3:19 acylsucrose but not F-type S3:22 acylsucrose. (**A**) LC-MS analysis of *in vitro* enzyme assay products indicates that SpASFF1 hydrolyzed P-type S3:19 (5^R2^,10^R3^,4^R4^) acylsucrose yielding two compounds with *m/z* = 533.3. This *m/z* is consistent with an acylglucose product with a G3:19 (4,5,10) configuration; the two peaks represent the α and β anomers of the acylglucose. (B) LC-MS analysis of *in vitro* assays with F-type S3:22 (12^R3^,5^R4^,5^R3’^) acylsucrose indicates no hydrolysis products with SpASFF1. Note: Acylglucose structure is inferred from collision induced dissociation-mediated fragmentation (Fig. S9).

## Discussion

The results described above show that *S. pennellii* LA0716 synthesizes acylglucoses from P-type acylsucroses via the action of a previously uncharacterized trichome invertase (Fig. 5A), a homolog of the most venerable enzyme in the history of biochemistry, yeast invertase. The canonical GH32 enzyme was first characterized in the 1840’s through studies of ‘optical inversion’ of cane sugar (sucrose) into a mixture of glucose and fructose. The enzyme was assayed two decades later, and its study by Maud Leonora Menten and Leonor Michaelis led to the theory of enzyme kinetics early in the 20^th^ Century (*31*). Since that time, other general metabolic activities were identified for diverse GH32 β-fructofuranosidases, including plant glycan biosynthesis, cell wall modification, and hormone metabolism (*43, 44*).

Our results stand in contrast to the previously proposed direct synthesis of acylglucoses from UDP-glucose and free fatty acids (*27–29,45*). Steffens and co-workers identified two glucosyltransferases and an SCPL acyltransferase from *S. pennellii* capable of generating *1-O-*mono- and 1,2-*O*-di-acylglucoses *in vitro* (*27–29,45*). Multiple lines of evidence indicate that these enzymes are not involved in *S. pennellii* acylglucose biosynthesis. First, acylglucoses generated *in vitro* by these enzymes are structurally distinct from the 2,3,4-*O*-tri-acylglucoses detected from *S. pennellii* (*19*); the *in vitro* products possessed two acyl chains instead of three and were acylated at position R1. Next, comparative transcriptomic data suggest that the SCPL acyltransferase shows similar expression levels in *S. lycopersicum* M82 and *S. pennellii* LA0716, yet there are no acylglucoses detected in M82 (*46*). Additionally, the SCPL acyltransferase described by Li et al. (2000) is encoded on chromosome 10 (Solyc10g049210), in a region not implicated in acylglucose accumulation in QTL mapping studies (*30*). In contrast, QTLs linked to acylglucose accumulation in *S. pennellii* on chromosomes 4 and 11 include *SpASAT2* and *SpASAT3*, suggesting a connection between acylsucrose and acylglucose biosynthesis (*23, 30*).

As the acylsucroses and acylglucoses in *S. pennellii* differ only by the presence or absence of a furanose ring (Fig. 1A), we hypothesized that a glycoside hydrolase converts the *S. pennellii* acylsucroses (*42*) to acylglucoses. Three glycoside hydrolase genes were identified in the third acylglucose-linked QTL on chromosome 3. These genes represent members of glycoside hydrolase (GH) families 32, 35, and 47 (Table S2). Most characterized plant GH35 enzymes act as β-galactosidases while GH47 enzymes function as α-mannosidases in post-translational protein modification (*47, 48*). Thus, these were not compelling candidates for cleavage of acylated sucrose substrates. Conversely, GH32 enzymes act on a variety of β-fructofuranosides in plants, including sucrose and fructans (*38, 43*). Our results indicate that SpASFF1 is a ‘derived’ β-fructofuranoside, with an active site that can accommodate pyranose-but not furanose-acylated sucrose esters. Understanding the structural features that allow SpASFF1 to hydrolyze P-type acylsucroses could inform engineering of novel specialized metabolites in plants and microbes.

We identified the GH32 SpASFF1 β-fructofuranosidase as being necessary and sufficient for conversion of P-type acylsucroses into acylglucoses. The most direct evidence is that ablation of the *SpASFF1* gene using CRISPR gene editing led to acylsucrose-accumulating wild tomato *S. pennellii* LA0716 mutants, showing that the enzyme is necessary for production of acylglucoses (Fig. 3). Multiple lines of genetic and biochemical evidence support the hypothesis that SpASFF1 uses P-type acylsucrose substrates. For example, no acylglucoses were detected in the F-type acylsucrose-producing introgression line IL3-5, despite the presence of *SpASFF1* in the introgressed region (Fig. 4A, and Fig. S1). In contrast, transgenic trichome expression of the SpASFF1 invertase in the P-type acylsucrose-producing SpASAT2 and SpASAT3 double introgression line *S. lycopersicum* BIL6180 resulted in acylglucose accumulation (Fig. 4). Our *in vitro* assay results support the *in vivo* evidence that P-type acylsucroses are SpASFF1 substrates. *In vitro* assays with recombinant SpASFF1 demonstrated conversion of the purified P-type S3:19 (5^R2^,10^R3^,4^R4^) to the cognate acylglucose G3:19 (4,5,10) (Fig. 5A). In contrast, the enzyme was inactive against F-type S3:22 (12^R3^,5^R4^,5^R3’^) (Fig. 5B) and did not hydrolyze unacylated sucrose (Fig. S10).

Taken together, these data indicate that *S. pennellii* acylglucose metabolism results from evolution of a three-gene epistatic system, where the innovation of P-type acylsucrose synthesis by modification of the core BAHD acyltransferases potentiated evolution of SpASFF1 to produce acylglucoses. Our results reveal that a member of the GH32 β-fructofuranosidase enzyme family acquired expression in the trichome glandular tip cell (Fig. 2) and the ability to cleave acylated sucrose (Fig. 5) to increase the diversity of Solanaceae trichome specialized metabolites. This is a remarkable evolutionary innovation, where a member of an enzyme family long recognized as important in general metabolism was co-opted into specialized metabolism by the ‘blind watchmaker’ of evolution.

Acylsugar accumulation is widespread throughout the Solanaceae with occurrences in genera as distantly related as *Salpiglossis* and *Solanum*, sharing a last common ancestor > 30 Mya (*8, 10, 49*). While acylsugars show wide structural variation in the number and length of acyl chains throughout the family, sucrose is the most prominent sugar core. Acylsucroses accumulate in genera whose lineages diverged < 20 Mya, such as *Solanum* and *Physalis* (*8, 21*) but also accumulate in species representing earlier diverging lineages, including *Salpiglossis* and *Petunia* (*10, 12*). In addition, acyl chains are present on the furanose ring in at least some members of each of these genera, suggesting that accumulation of F-type acylsucroses evolved long ago.

Though apparently limited in distribution relative to acylsucroses, acylglucoses occur in diverse genera including *Solanum, Datura*, and *Nicotiana* (*16, 19, 20*). Despite the fact that acylglucose accumulation is common to species in both *Solanum* and *Nicotiana* – which diverged approximately 24 Mya – the differences in *SpASFF1* gene expression and SpASAT substrate specificity that facilitated acylglucose accumulation in *S. pennellii* arose in the ~7 million years since divergence from the last ancestor in common with *S. lycopersicum* (*11, 49, 50*). This supports independent evolutionary origins of acylglucoses in distinct lineages. In the *Solanum* genus, P-type acylsucroses are a pre-requisite for acylglucose accumulation. The predominance of F-type acylsucroses within the Solanaceae may explain the relative rarity of acylglucoses in the family. However, characterization of the ASAT enzymes responsible for acylsucrose biosynthesis in *Salpiglossis, Petunia*, and *Solanum* demonstrates multiple changes in enzyme substrate specificity throughout the evolutionary history of the acylsucrose pathway (*10–12*). Plasticity of the acylsugar pathway may have given rise to P-type acylsucroses multiple times throughout evolutionary history. If so, this would provide independent opportunities for co-option of acylglucose-producing glycoside hydrolases into the acylsugar pathway. Are the enzymes responsible for hydrolyzing acylsucroses to yield acylglucoses restricted to the GH32 family or have other enzyme families evolved in different acylglucose-accumulating lineages? If and to what extent multiple origins of acylglucose biosynthesis share common features remains to be explored.

Over the past decade, discovery of pathways and enzymes of plant specialized metabolism has improved at an increasing rate. Prior to this time, taxonomic restriction of specialized metabolism biased deep analysis towards pathways found in model organisms: for instance, glucosinolates in Arabidopsis, cyclic hydroxamic acids in maize and other well-studied grasses and isoflavonoids in Medicago and soybean. Dramatic improvements in sensitivity and selectivity of MS- and NMR-based analytical chemistry helped broaden the scope of well-studied metabolic networks (*51, 52*). In parallel, development of species-agnostic DNA sequencing and functional genomics screening tools (notably, viral induced gene silencing and genetic transformation), permitted rigorous correlation of *in vitro* activities and *in vivo* phenotypes. The rapid advancement of gene editing techniques using CRISPR-Cas on agriculturally-important and undomesticated species dramatically expands the specialized metabolism functional genomics toolkit. Not only do these methods allow direct tests of *in vivo* function, but also allows elimination of the t-DNA by simple genetic crossing. The removal of the T-DNA permits growing edited mutants in agricultural fields or common gardens with lower regulatory barriers. For example, the *spasff1* mutant lines can be used to understand the impacts of acylsucroses versus acylglucoses on the fitness of *S. pennellii* both in the greenhouse and field. Such studies could lead to crops with novel natural pesticides, broaden our understanding of the roles of specialized metabolites in mediating environmental interactions, and inform our understanding of the mechanisms underpinning specialized metabolic evolution.

## Materials and Methods

### Plant material

Seeds of *S. lycopersicum* M82 were obtained from the C.M. Rick Tomato Genetics Resource Center (TGRC; University of California, Davis, CA); seeds of IL3-5, BIL6180, and BIL6521 were obtained from Dr. Dani Zamir (Hebrew University of Jerusalem, Rehovot, Israel) (*36*); seeds of *S. pennellii* LA0716 were generously provided by Dr. Martha Mutschler (Cornell University, Ithaca, NY). Young plants were grown in 9-cm pots in peat-based propagation mix (SunGro, Agawam, MA). *S. lycopersicum* and introgression lines were watered four times weekly and supplemented once weekly with ½ strength Hoagland’s solution; *S. pennellii* was watered once weekly and supplemented once weekly with ½ strength Hoagland’s solution. Plants used for analysis were grown in a growth chamber under a 16-h photoperiod (190 μmol m^−2^ s^−1^ photosynthetic photon flux density (PPFD)) with 28°C day and 22°C night temperatures set to 50% relative humidity. BIL lines used for crosses were grown in a soil mix consisting of four parts SureMix (Michigan Grower Products, Inc., Galesburg, MI) to one part sand in a greenhouse with a daytime maximum temperature of 30° C and a nighttime minimum temperature of 16° C; sunlight was supplemented with high pressure sodium bulbs on a 16/8 light/dark cycle. For seed production, *S. pennellii asff1* T_0_ plants were grown in soil containing one part Canadian sphagnum (Mosser Lee Co., Millston, WI), one part coarse sand (Quikrete, Atlanta, GA), one part white pumice (Everwood Farm, Brooks, OR), and one part redwood bark (Sequoia Bark Sales, Reedley, CA) supplemented with 1.8 kg crushed oyster shell (Down to Earth Distributors Inc., Eugene, OR), 1.8 kg hydrated lime (Bonide Products, Inc., Oriskany, NY), and 0.6 kg triple super phosphate (T and N Inc., Foristell, MO) per cubic meter.

### Acylsugar analysis

Leaf surface acylsugars were extracted from single leaflets with 1 mL of a mixture of isopropanol (J.T. Baker, Phillipsburg, NJ):acetonitrile (Sigma-Aldrich, St. Louis, MO):water (3:3:2) with 0.1% formic acid and 1 μM telmisartan (Sigma-Aldrich, St. Louis, MO) as an HPLC standard. The leaf tissue was gently agitated on a rocker in this extraction solvent for 2 min. The extraction solvent was collected and stored in 2 mL LC-MS vials at −80°C.

LC-MS samples (both enzyme assays and plant samples) were run on a Waters Acquity UPLC coupled to a Waters Xevo G2-XS QToF mass spectrometer. 10 μL of the acylsugar extracts were injected into an Ascentis Express C18 HPLC column (10 cm × 2.1 mm, 2.7 μm) (Sigma-Aldrich, St. Louis, MO), which was maintained at 40°C. The LC-MS methods used the following solvents: 10 mM ammonium formate, pH 2.8 as solvent A, and 100% acetonitrile as solvent B. Compounds were eluted using one of two gradients.

A 7-min linear elution gradient consisted of 5% B at 0 min, 60% B at 1 min, 100% B at 5 min, held at 100% B until 6 min, 5% B at 6.01 min and held at 5% until 7 min. A 21-min linear elution gradient consisted of 5% B at 0 min, 60% B at 3 min, 100% B at 15 min, held at 100% B until 18 min, 5% B at 18.01 min and held at 5% B until 21 min.

The MS settings were as follows for negative ion-mode electrospray ionization: 2.00 kV capillary voltage, 100°C source temperature, 350°C desolvation temperature, 600 liters/h desolvation nitrogen gas flow rate, 35V cone voltage, mass range of m/z 50 to 1000 with spectra accumulated at 0.1 seconds/function. Three separate acquisition functions were set up to test different collision energies (0V, 15V, 35V).

The MS settings were as follows for positive ion-mode electrospray ionization: 3.00 kV capillary voltage, 100°C source temperature, 350°C desolvation temperature, 600 liters/h desolvation nitrogen gas flow rate, 35V cone voltage, mass range of m/z 50 to 1000 with spectra accumulated at 0.1 seconds/function. Three separate acquisition functions were set up to test different collision energies (0V, 15V, 45V). Lockmass correction was performed using leucine encephalin as the reference compound for data acquired in both negative and positive ion mode.

### Acylsugar quantification

To accurately quantify total acylsugars, samples were saponified before LC-MS analysis and sugar cores quantified with authentic isotopically labelled standards. A leaflet was immersed in 2 mL dichloromethane (VWR International, Radnor, PA) and 500 μL water with 30 s vortexing. After phase separation, 1 mL of the dichloromethane layer was removed to a borosilicate glass vial and evaporated to dryness under flowing air. Dried samples were dissolved in 1 mL acetonitrile with 0.1% formic acid for storage. 20 μL aliquots of acylsugar extracts were dried in 1.7 mL microcentrifuge tube using a SpeedVac and dissolved in 100 μL methanol. An equal volume of 3 N aqueous ammonia solution (Sigma-Aldrich, St. Louis, MO) was added, and the reaction was incubated in a sealed 1.5 mL microcentrifuge tube for 48 hrs in a fume hood. Before LC-MS analysis, samples were dissolved in 200 μL ammonium bicarbonate (pH 7-8) in 90% acetonitrile containing 0.5 μM ^13^C_12_-sucrose and 0.5 μM ^13^C_6_-glucose as internal standards and transferred to a 2 mL LC-MS vials. Compounds were analyzed on a *Waters ACQUITY TQD* Tandem Quadrupole UPLC/MS/MS system (Waters, Milford, MA). Ten microliters of the acylsugar extracts were injected into a Waters ACQUITY UPLC BEH amide column (2.1×100 mm, 1.7 μM), in a column oven with temperature of 40°C with flow rate of 0.5 mL/min. The LC-MS methods used 10 mM ammonium bicarbonate pH 8 in 50% acetonitrile as Solvent A and 10 mM ammonium bicarbonate pH 8 in 90% acetonitrile as Solvent B. The chromatography gradient was: 100% B at 0 min, 0% B at 5 min, 100% B at 5.01 min, held at 100% B until 10 min. Multiple-reaction monitoring (MRM) mode was operated to detect each sugar. For glucose: parent ion, *m/z* 179; product ion, *m/z* 89; cone voltage, 16V; collision energy, 10V. For ^13^C_6_-glucose: parent ion, *m/z* 185; product ion, *m/z* 92; cone voltage, 16V; collision energy, 10V. For sucrose: parent ion, *m/z* 341; product ion, *m/z* 89; cone voltage, 40V; collision energy, 22V. For ^13^C_12_-sucrose: parent ion, *m/z* 353; product ion, *m/z* 92; cone voltage, 40V; collision energy, 22V. Quantification of glucose and sucrose were conducted by standard curves with authentic glucose and sucrose standards (Sigma-Aldrich, St. Louis, MO).

### Acylsucrose purification

All purifications were performed using a Waters 2795 Separations Module (Waters, Milford, MA) and an Acclaim 120 C18 HPLC column (4.6 × 150 mm, 5 μm; Thermo Scientific, Waltham, MA) with a column oven temperature of 30°C and flow rate of 1 mL/min. The mobile phase consisted of water (Solvent A) and acetonitrile (Solvent B). Fractions were collected using a 2211 Superrac fraction collector (LKB Bromma, Stockholm, Sweden).

For purification of acylsucroses from *S. pennellii* LA0716, approximately 75 g fresh above-ground tissue of mature *S. pennellii asff1-1* was harvested into a 1 L glass beaker to which 500 mL 100% methanol was added. Tissue was stirred for 2 minutes and filtered through Miracloth (EMD Millipore, Billerica, MA) pre-wetted with methanol into a 1-L round bottom flask. Solvent was removed with a rotary evaporator in a water bath held between 35 and 40° C and the residue dissolved in 5 mL acetonitrile. A 5-μL aliquot of this solution was diluted 1000-fold in 9:1 water/acetonitrile with 0.1% formic acid for chromatographic purification. The S3: 19 compound was purified from 20 injections of 100 μL each using a linear elution gradient of 1% B at 0 min, 63% B at 10 min, 65% B at 30 min, 100% B at 35 min brought back to 1% B at 35.01 min and held at 1% B until 40 min. Eluted compounds were collected in 10 second fractions. Fraction collection tubes contained 333 μL 0.1% formic acid in water, and the S3: 19 product eluted at 18–19 min.

For purification of the S3:22 acylsucrose from *S. lycopersicum* M82, approximately 75 g fresh above-ground tissue was harvested from mature plants into a 500-mL glass beaker to which 250 mL 100% methanol was added. The tissue was stirred for 2 min and filtered through Miracloth into a 1 L round-bottom flask. Methanol was removed with a rotary evaporator and residue dissolved in 5 mL acetonitrile. This solution was diluted 50-fold in 9:1 water/acetonitrile with 0.1% formic acid for further processing. The S3:22 compound was purified from 10 injections of 100 μL using a linear elution gradient of 1% B at 0 min, 50% B at 5 min, 70% B at 30 min, 100% B at 32 min and held at 100% until 35 min brought back to 1% B at 35.01 min and held at 1% B until 40 min. Eluted compounds were collected in 1 minute fractions. Fraction collection tubes contained 333 μL 0.1% formic acid in water, and the S3:19 product eluted at 7–8 minutes.

### qPCR analysis

Tissue of 10-week-old *S. pennellii* LA0716 and *S. lycopersicum* M82 were harvested, with stems flash-frozen in liquid nitrogen and trichomes shaved into 1.5 mL microcentrifuge tubes with a clean razor blade. Trichomes and denuded stems were kept in liquid nitrogen and ground with plastic micropestles in 1.5 mL microcentrifuge tubes. RNA was extracted from ground trichomes and stems (six biological replicates for each species and tissue type) using the RNeasy Plant Mini Kit (Qiagen, Hilden, Germany) according to the manufacturer’s instructions. For each sample, 250 ng of RNA as quantified using a Nanodrop 2000c (Thermo Fisher Scientific, Waltham, MA) was used to synthesize cDNA using SuperScript III reverse transcriptase (Invitrogen, Carlsbad, CA). qRT-PCR was carried out using SYBR Green PCR Master Mix on a QuantStudio 7 Flex Real-Time PCR System (Applied Biosystems, Warrington, UK) using the following cycling conditions: 48°C for 30 min, 95°C for 10 min, 40 cycles of 95°C for 15 s and 60°C for 1 min followed by melt curve analysis. RT_ASFF_F and RT_ASFF_R primers were used to detect ASFF1 transcript; RT_EF-1a_F/R, RT_actin_F/R, and RT_ ubiquitin_F/R primers were used to detect transcripts of the *EF-1α, actin*, and *ubiquitin* genes, respectively (Table S3). For each biological replicate, relative levels of ASFF1 transcript were determined using the ΔΔC_t_ method (*53*) and normalized to the geometric mean of EF-1α, actin, and ubiquitin transcript levels.

### Genotyping of progeny of BIL6521 x BIL6180

DNA from the progeny of the cross between BIL6521 and BIL6180 was extracted from leaf material that were archived on FTA PlantSaver cards (GE Healthcare, Uppsala, Sweden) and purified according to the manufacturer’s specifications. Extracted DNA from the FTA cards were used for PCR amplification with GoTaq green mastermix to genotype the sample using 04g011460_Marker_Indel-F/R and ASFF_Chr3_Indel_002_F/R (Table S3).

### DNA construct assembly

All Sanger DNA sequencing confirmation in this study was performed with the indicated sequencing primers at the Research Technology Support Facility Genomics Core, Michigan State University, East Lansing, MI.

For *proSpASFF1::SpASFF1* ORF – (pK7WG), a 1.8 kb region of the upstream region and open reading frame of *SpASFF1* was split into four amplicons using four sets of primers: ASFF_001_F/R, ASFF 002_F/R, ASFF_003_F/R, and ASFF_004_F/R (Table S3). The first and fourth amplicon contained adapters for assembly into pENTR-D-TOPO that has been digested with NotI/AscI respectively. The construct was assembled using NEB Gibson assembly according to manufacturer specifications (NEB, Ipswich, MA). The construct was verified by Sanger sequencing using M13 Forward and T7 promoter primers. The insert was subcloned into pK7WG (*54*) using LR clonase II enzyme mix (Thermo Scientific, Waltham, MA) according to manufacturer instructions. Presence of the insert was determined by colony PCR using ASFF_001F/R. Completed vectors were transformed into *Agrobacterium* strain AGL0. Leaf material from recovered plants were archived on FTA PlantSaver cards (GE Healthcare, Uppsala, Sweden) and genotyped by PCR amplification with GoTaq green mastermix and pK7WG-Kan-F/R primers (Table S3)

For *proSpASFF1*::GFP/GUS – (pKGWFS7), a 1.8 kb region of the upstream region of ASFF1 was amplified from *S. pennellii* LA0716 genomic DNA using the primers ASFF_promoter_F1/R1 (Table S3). pENTR-D-TOPO was digested with NotI/AscI to linearize the vector and create overhangs compatible for Gibson assembly. The amplicon also contained adapters for insertion into pENTR-D-TOPO digested with NotI/AscI. Constructs were Sanger sequenced using M13F/R primers in addition to the ASFF_promoter_F1/R1 primers. LR clonase II mix was used to subclone the fragment into pKGWFS7 (*54*). Construct was transformed into Agl0 for plant transformation using the described protocol.

The CRISPR-ASFF1 vector was constructed as follows. CRISPR sgRNAs were designed using the site finder toolset in Geneious^®^ v10 (www.geneious.com). Two target sequences located on the exon were selected for their high on-target activity scores, based on a published algorithm (*55*), and low off-target scores against published *S. pennellii* genome database (*56*). Each sgRNA was obtained as a gBlock synthesized *in vitro* by IDT (www.idtdna.com) (Table S3) and subsequently assembled with pICH47742::2×35S-5’UTR-hCas9(STOP)-NOST (Addgene #49771, kindly provided by Dr. Sophien Kamoun, Sainsbury Lab, Norwich, UK) (*57*), pICH41780 (Addgene plasmid # 48019) and pAGM4723 (Addgene plasmid # 48015, both gifts from Dr. Sylvestre Marillonnet) (*58*) and pICSL11024 (Addgene plasmid # 51144, a gift from Dr. Jonathan D. Jones, Sainsbury Lab, Norwich, UK) using Golden Gate Assembly. In short, the restriction-ligation reactions (20 μL) were set up by mixing 15 ng of synthesized sgRNAs with 1.5 μL T4 ligase buffer (NEB), 320 U of T4 DNA ligase (NEB), 1.5 μL BSA (0.1 mg/mL, NEB), 8 U of *Bpi*I (Thermoscientific) and 100–200 ng of the intact plasmids. The reactions were incubated at 37°C for 30s, followed by 26 cycles (37°C, 3 minutes; 16°C, 4 minutes) and then incubated at 50°C for 5 minutes and 65°C for 5 minutes. The ligated products were directly used to transform *E. coli* competent cells. Positive clones were chosen based on colony PCR and sequenced at the MSU RTSF facility using the pAGM4723_SeqF1, pAGM4723_SeqR1, pICSL11024_SeqF1, pICH47742CAS9_SeqF2, pICH47742_SeqF1, pICH41780_SeqR1, and ASFF_SeqR primers (Table S3). The construct was transformed into *S. pennellii* LA0716 using the plant transformation protocol described below. Leaf material from recovered plants were archived on FTA PlantSaver cards (GE Healthcare, Uppsala, Sweden) and genotyped by PCR amplification with ASFF_F/R, followed by Sanger sequencing with ASFF_SeqR (Table S3).

For *spasff1* line transcript analysis, RNA was extracted from *spasff1-1-1/1-1-2* lines using RNeasy plant mini kit according to the kit specifications (Qiagen, Venlo, Netherlands). RNA was quantified using a Nanodrop 2000c (Thermofisher, Waltham, MA). 1 μg of RNA was used for cDNA synthesis using Superscript II Reverse Transcriptase according to the manufacturer’s specifications. The primers, ASFF1_transcript_amp_01F/R (final concentration: 0.5 μM), were used to amplify the region within the *ASFF1* CDS, which was cloned into pMINI-T 2.0 (NEB, Ipswich, MA). T7 and SP6 promoter primers were used for Sanger sequence confirmation of the inserts, and ClustalW was used for alignment of the transcripts (https://www.ebi.ac.uk/Tools/msa/clustalo/).

### Competent cell preparation and transformation of constructs into Agrobacterium

A single colony of AGL0 or LBA4404 *Agrobacterium* was inoculated into two 5 mL cultures of YEP media (10 g yeast extract, 10 g Bacto peptone, and 5 g NaCl per liter, pH 7) with Rifampicin (50 μg/mL). Cultures were incubated in 18×150 mm borosilicate glass test tubes with foam plugs overnight at 30°C, shaking at 200 rpm. 190 mL of LB was inoculated with 10 mL of the overnight cultures in an autoclaved 500 mL Erlenmeyer flask. Cultures were grown in a shaking incubator (200 rpm) at 30°C to OD_600_ = 1.0, incubated on ice for 10 minutes, and centrifuged at 4°C in 50 mL conical tubes at 3,200g for 5 minutes. Pellets were resuspended in 1 mL of sterile 20 mM CaCl_2_ and 100 μL aliquots, dispensed into sterile, pre-chilled 1.7 mL microcentrifuge tubes, snap frozen using liquid nitrogen and stored at −80°C

For *Agrobacterium* transformation, 1 μg of construct DNA purified using an Omega EZNA plasmid DNA mini kit I (Omega Bio-Tek, Norcross, GA) as added to the frozen *Agrobacterium* aliquots on ice. Cells were thawed in a 37°C water bath for 5 minutes, mixed well by flicking and snap frozen in liquid nitrogen. Cells were thawed, and 1 mL of YEP added to the tube. The transformations were incubated at 28°C and 200 rpm for 4 hours. Cells were centrifuged at 17,000g for 30 seconds, the supernatant decanted and the cell pellet resuspended in 100 μL of fresh YEP. The cell pellet was resuspended and the entire suspension was plated onto an LB plate with Spectinomycin (100 μg/mL).

Presence of the insert containing vector was verified by colony PCR. Colonies were collected with a pipette tip and resuspended in 20 μL of sterile water. 2 μL of the cell suspension was added to a PCR tube with a reaction with a final volume of 25 μL. GoTaq green mastermix (2x) was used for the colony PCR according to the manufacturer’s specifications (Promega, Madison, WI). Primers (0.4 μM final concentration) pertaining to the insert were used for amplification.

### Plant transformation

In all cases, petri plates containing plant tissue were sealed with a single layer of micropore paper tape (3M, Maplewood, MN).

Transformation of *S. lycopersicum* and *S. pennellii* LA0716 was performed using AGL0 using a modification of published protocols (*59, 60*). 50–60 seeds were incubated in 40% bleach, agitating for 5 minutes. Seeds were rinsed six times, each with 40 mL of sterile ddH_2_O with 5 minutes of rocking and decanting of wash solution. A flame-sterilized spatula was used to distribute the seeds onto the surface of 1/2x MSO medium (*59*) in a PhytaTray II (Sigma-Aldrich, St. Louis, MO). Containers were incubated at 25°C on a 16/8 light/dark cycle with a light intensity of 70 μmol m^−2^ s^−2^ PPFD.

At day eight for *S. lycopersicum* or day 11 for *S. pennellii* LA0716, the seedlings were removed from the 1/2x MSO medium jar. The hypocotyl and radicle were excised and discarded. The cotyledon explant was placed on a sterile petri dish. 1–2 mm was removed from the base and tip of the cotyledon. An autoclaved piece of Whatman #1 filter paper (GE Healthcare, Uppsala, Sweden) was placed on the surface of a sterile D1 media plate (*59*) on which the cotyledons were placed adaxial side up. Approximately 100 explants were added per plate. The plates were placed in the same conditions for two days until day 10.

For co-cultivation, the Agrobacterium containing the construct was streaked out onto an LB plate containing the appropriate antibiotic. A single colony was inoculated into a 25 mL LB culture with the same antibiotic plus Rifampicin (50 mg/L) in a 250 mL Erlenmeyer flask. The culture was incubated at 30°C in a shaking incubator (225 rpm) for 2 days. The culture was transferred to a sterile 50 mL conical tube and centrifuged at 3,200g for 10 minutes at 20°C. The supernatant fluid was decanted and 10 mL of MSO media (*59*) was added to the tube (with no pellet resuspension). The cell pellet was centrifuged at 2,000g for 5 minutes and this washing step was then repeated. The cell pellet was re-suspended in 10–20 mL of MSO liquid media. Absorbance of the culture was measured at 600 nm. The suspension was diluted with MSO to OD_600_ = 0.5. Acetosyringone dissolved in DMSO was added at a final concentration of 375 μM and 5 mL of the Agrobacterium suspension pipetted onto the cotyledons on the plate and incubated with swirling at room temperature for 10 minutes, at which point the excess culture was pipetted off. Using a scalpel, cotyledons were transferred to a fresh D1 medium plate containing autoclaved Whatman paper. Approximately 50 cotyledons per plate were placed abaxial side up. Plates were incubated at 24°C for 2 days with a 16/8 day-night cycle at 70 μmol m^−2^ s^−2^ PPFD.

For transgenic callus selection, two days after cocultivation, the cotyledons were transferred directly onto sterile 2Z media plates (*60*) containing 100 μg/mL kanamycin and 200 μg/mL timentin (no filter paper). Explants were placed abaxial side up with 20–30 cotyledons per plate. Plates were incubated at the same growth conditions for 10 days. Cotyledons were then transferred to a sterile Petri dish and, using a scalpel, calluses were cut and then placed onto fresh 2Z media plates with the same selection. Subsequently, explants were transferred to new 2Z plates every two weeks. Throughout the process, dying tissue was removed, and growing tissue was placed on the media. Five to eight weeks after cocultivation, shoots were harvested from the explants and placed into Phytatray II (Sigma-Aldrich, St. Louis, MO) containing 100 mL of MSSV media (*60*) supplemented with Timentin (100 μg/mL), Kanamycin (50 μg/mL), and Indole-3 butyric acid (1 μg/mL). MSSV-containing Phytatrays were incubated at the same growth conditions (16/8 at 70 μmol m^−2^ s^−2^ PPFD). Shoots were monitored for leaf and root production, and shoots with roots and leaves were placed into pots containing RediEarth soil. Flats were covered with a plastic dome in the same growth conditions. Domes were removed from flats after three to four days.

### Transient expression and purification of SpASFF1 protein

The ASFF1 CDS was amplified from *S. pennellii* LA0716 trichome cDNA using the ASFF_F and ASFF_R primers (Table S3) and cloned into the pGEM backbone using the pGEM-T Easy cloning kit (Promega, Madison, WI). The ASFF1 CDS was subsequently re-amplified with the (pEAQ-HT)-ASFF-His_F and (pEAQ-HT)-ASFF-His_R primers (Table S3) to add adapters for Gibson assembly. The resulting PCR product was transferred to pEAQ-HT vector (*41*) previously digested with NruI-HF and SmaI restriction enzymes (New England Biolabs, Ipswich, MA) using 2x Gibson Assembly master mix (New England Biolabs, Ipswich, MA) according to the manufacturer’s instructions to create an expression clone coding for the full-length protein with a C-terminal 6x His tag (ASFF1-HT-pEAQ). The completed vector was subsequently transformed into LBA4404 cells as described above. For transient expression, an *A. tumefaciens* LBA4404 strain carrying the ASFF1-HT-pEAQ construct was streaked onto LB agar containing 50 μg/mL rifampicin and 50 μg/mL kanamycin and incubated for 3 days at 28°C. Single colonies were used to inoculate 250-mL Erlenmeyer flasks containing 50 mL YEP medium with 50 μg/mL rifampicin and 50 μg/mL kanamycin; cultures were incubated at 28°C and 300 rpm overnight. Cultures were harvested by centrifugation at 800g and 20°C for 20 min. Supernatant was discarded and the resulting loose pellet resuspended in 50 mL of buffer A (10 mM 2-ethanesulfonic acid (MES; Sigma-Aldrich, St. Louis, MO) pH 5.6, 10 mM MgCl_2_). This cell suspension was centrifuged at 800g and 20°C for 20 min and the resulting pellet resuspended to a final OD_600_ = 1.0 with buffer A. A 200 mM solution of acetosyringone (Sigma-Aldrich, St. Louis, MO) dissolved in DMSO was added to the suspension at a final concentration of 200 μM and the suspension incubated at room temperature with gentle rocking for 4 h. This suspension was infiltrated into fully expanded leaves of six-week-old *Nicotiana benthamiana* plants using a needle-less 1-mL tuberculin syringe. Plants were grown under 16-h photoperiod (70 μmol m^−2^ s^−1^ PPFD) and constant 22°C set to 70% relative humidity. At 8 days post-infiltration, 28 g infiltrated leaves were harvested, de-veined, and flash-frozen in liquid nitrogen. Tissue was powdered under liquid nitrogen with mortar and pestle and added to 140 mL ice-cold buffer B (25 mM 3-[4-(2-hydroxyethyl)piperazin-1-yl]propane-1-sulfonic acid (EPPS) pH 8.0, 1.5 M NaCl, 1 mM ethylenediaminetetraacetic acid (EDTA) with 2 mM dithiothreitol (DTT), 1 mM benzamidine, 0.1 mM phenylmethansesulfonylfluoride (PMSF), 10 μM *trans*-epoxysuccinyl-L-leucylamido(4-guanidino)butane (E-64), and 5% (w/v) polyvinylpolypyrrolidone (PVPP); all reagents were obtained from Sigma-Aldrich, St. Louis, MO except DTT obtained from Roche Diagnostics, Risch-Rotkreuz, Switzerland). The mixture was stirred for 4 h at 4°C, filtered through six layers of Miracloth and centrifuged at 27,000g, 4°C for 30 min. The supernatant was decanted and passed through a 0.22 μm polyethersulfone filter (EMD Millipore, Billerica, MA) before being loaded onto a HisTrap HP 1 mL affinity column and eluted using a gradient of 10 to 500 mM imidazole in buffer B using an ÄKTA start FPLC module (GE Healthcare, Uppsala, Sweden). Fractions were analyzed by SDS-PAGE and the presence of ASFF1-HT confirmed by immunoblot using the BMG-His-1 monoclonal antibody (Roche, Mannheim, Germany) to detect His-tagged proteins. Purified ASFF1-HT was subsequently transferred to 100 mM sodium acetate pH 4.5, 50% glycerol using a 10DG desalting column (Bio-Rad, Hercules, CA). Protein was quantified against a standard curve of bovine serum albumin (Thermo Fisher Scientific, Waltham, MA) using a modified Bradford reagent (Bio-Rad, Hercules, CA) according to the manufacturer’s instructions.

### Enzyme assays

For activity assays, 100 ng ASFF1-HT or 1 μg *Saccharomyces cerevisiae* invertase (Cat. No. I4504, Grade VII, Sigma-Aldrich, St. Louis, MO) and 0.1 nmol F- or P-type acylsucrose or 10 nmol sucrose (Sigma-Aldrich, St. Louis, MO) were added to 30 μL 50 mM sodium acetate, pH 4.5 in 250-μL thin-wall PCR tubes. Reactions were incubated for 1 h at 30°C and stopped by addition of 60 μL 1:1 acetonitrile/isopropanol containing 1.5 μM telmisartan as internal standard and centrifuged 10 min at 16,000g to remove precipitated protein. The supernatant was transferred to 2 mL autosampler vials with 250-μL glass inserts and analyzed by LC-MS as described above.

### Statistical Analysis

All statistical analysis was performed using the ‘stats’ R package (R Core Team, 2017). One-way analysis of variance (ANOVA) was executed on acylsugar and transcript abundance data using the “aov” command. Between- and within-group variances were determined using the sum-of-squares values obtained from ANOVA; these values were subsequently used to determine the power of the ANOVA using the “power.anova.test” function. A post-hoc analysis by Tukey’s honestly significant difference (HSD) mean-separation test was executed using the “TukeyHSD” command with the results of one-way ANOVA as input. Implementation of these functions in the R console was as follows:

\# Object “Abundance” is a data table containing normalized relative quantities (“NRQ”) of ASFF1 transcript from various tissue types (“Tissue”): S. lycopersicum M82 trichomes (M82t), S. lycopersicum M82 stems (M82s), S. pennellii LA0716 trichomes (716t), and S. pennellii LA0716 stems (716s).

> Abundance

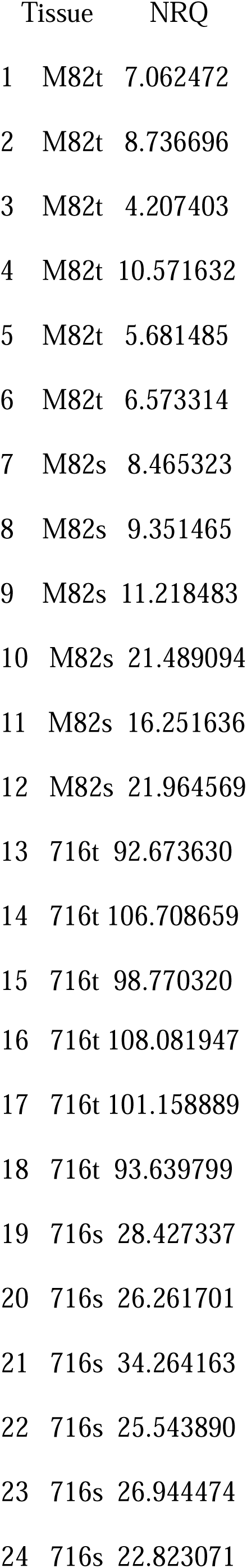

> asff1 <-aov(formula = NRQ ~ Tissue, data = Abundance)

> summary(asff1)

Df Sum Sq Mean Sq F value Pr(>F)

Tissue 3 32806 10935 448.7 <2e-16 ***

Residuals 20 487 24

---

Signif. codes: 0 ‘***’ 0.001 ‘**’ 0.01 ‘*’ 0.05 ‘.’ 0.1 ‘ ’ 1

\# Power is calculated to determine whether conclusions drawn using the chosen *p*-value are reliable. This step does not affect the results obtained by ANOVA or subsequent post-hoc analysis by Tukey’s HSD test. Between-group and within-group variances (“between.var” and “within.var”, respectively) can be manually calculated using values reported in the output of “summary”.

> power.anova.test(groups = 4, n = 6, between.var = 10935, within.var = 24.35, sig.level = 0.001)

Balanced one-way analysis of variance power calculation

groups = 4

n = 6

between.var = 10935

within.var = 24.35

sig.level = 0.001

power = 1

NOTE: n is number in each group

> TukeyHSD(asff1, conf.level = 0.95)

Tukey multiple comparisons of means

95% family-wise confidence level

Fit: aov(formula = NRQ ~ Tissue, data = Abundance)

$Tissue

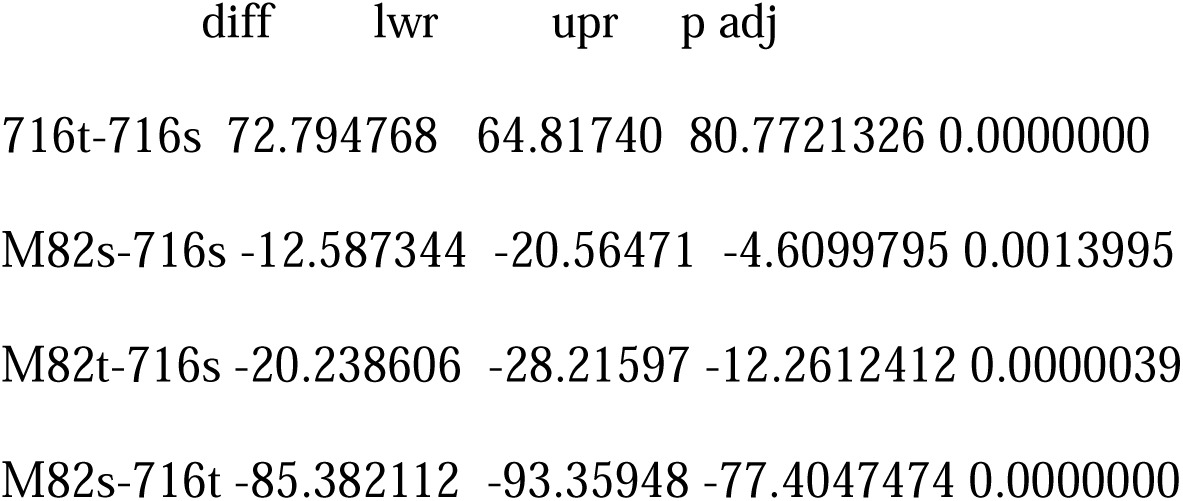

## Acknowledgments

### General

We thank Martha Mutschler, Dani Zamir, and the C. M. Rick Tomato Genetics Resource Center (UC Davis, CA) for providing germplasm essential to this study. Plant transformation was performed by Kathleen Imre. We also are grateful to Roger Chetelat and Xiaoqiong Qin for helpful advice about *S. pennellii* transformation; Daniel Jones and Lijun Chen and the MSU RTSF Mass Spectrometry and Metabolomics facility for analytical chemistry advice. We thank Eran Pichersky and members of the MSU Solanaceae Specialized Metabolism Evolution group for their advice.

### Funding

This work was funded by National Science Foundation grant IOS-PGRP-1546617 to RLL and National Institute of General Medical Sciences of the National Institutes of Health graduate training grant no. T32–GM110523 to BJL and DL.

### Author contributions

Design: BJL, DL, Y-RL, PF, ALS, RLL; Experiments and data analysis: BJL, DL, Y-LR; Interpretation and preparation of figures and table: BJL, DL, Y-RL, PF, ALS, RLL; Writing: BJL, DL, Y-RL, PF, ALS, RLL.

### Competing interests

No competing interests.

### Data and materials availability

Data that supports the findings of this study are available from the corresponding author upon reasonable request. Sequence data for *SpASFF*1 is available in Genbank: *SpASFF1* (MK297330). The following materials require an MTA: pEAQ-HT, pK7WG, pKGWFS7, pICH47742::2×35S-5’UTR-hCas9(STOP)-NOST, pICH41780, pAGM4723, and pICSL11024

