## Supplemental Figures and Tables for "Evolution of metabolic novelty: a trichome-expressed invertase creates specialized metabolic diversity in wild tomato"

**Supplementary materials**

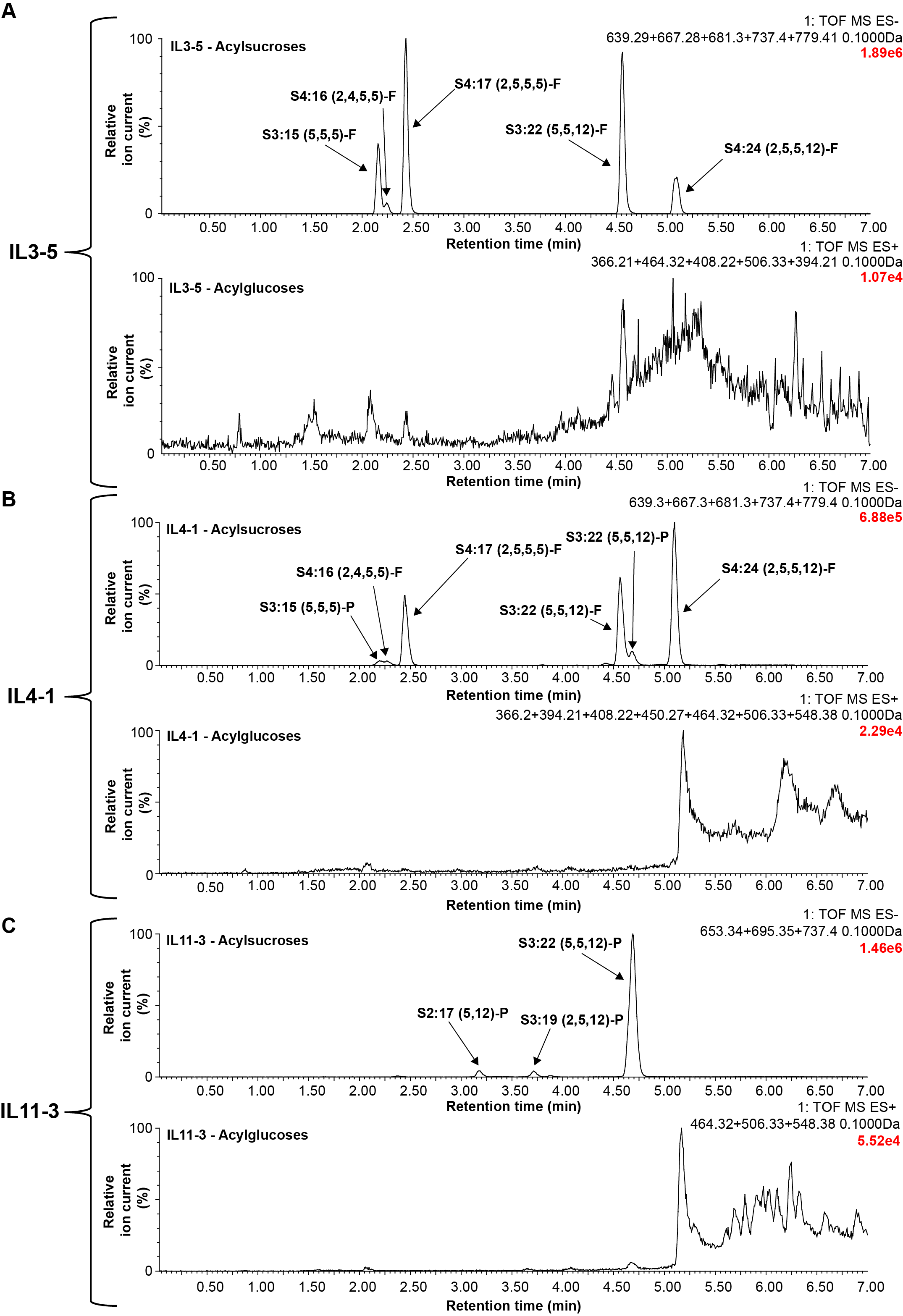

**Fig. S1. Acylglucoses are not detected in trichome extracts of IL3-5, 4-1, or IL11-3.**

LC-MS analysis using ESI- and ESI+ mode was used to detect acylsucroses and acylglucoses,respectively. Extracted ion chromatograms for ammonium adducts of expected possible acylglucoses showed no detectable peaks in any of the tested introgression lines (IL3-5, IL4-1, or IL11-3). The major acylsucroses identified were as follows: (**A**) For IL3-5, the major acylsugars identified were formate adducts of S3:15 (*m/z*: 639.28), S4:16 (*m/z*: 667.28), S4:17 (*m/z*: 681.30), S3:22 (*m/z*: 737.40), and S4:24 (*m/z*: 779.41). (**B**) For IL4-1, the major acylsugars present that were previously identified are: S3:15 (5,5,5)-P (*m/z*: 639.28), S4:16 (2,4,5,5) (*m/z*: 667.28), S4:17 (2,5,5,5) (*m/z*: 681.30), S3:22 (5,5,12) (*m/z*: 737.40), S3:22 (5,5,12)-P (*m/z*:737.40), and S4:24 (2,5,5,12) (*m/z*: 779.41). (**C**) For IL11-3, the major acylsugars present that were previously identified are: S2:17 (5,12) (*m/z*: 653.34), S3:19 (2,5,12) (*m/z*: 695.35), and S3:22 (5,5,12) (*m/z*:737.40). Note: The possible acylglucoses masses depended on the acylsucroses present, but all acylglucoses searched for were: G2:10 (5,5) (*m/z*: 366.21), G3:11 (2,4,5) (*m/z*: 394.21), G3:12 (2,5,5) (*m/z*: 408.22), G2:17 (5,12) (*m/z*: 464.32), G3:19 (2,5,12) (*m/z*: 506.33), G3:15 (5,5,5) (*m/z*: 450.27), G3:22 (5,5,12) (*m/z*: 548.38). The 7 min method was used for this LC-MS analysis, which is described in the Methods (mass window: 0.1Da). Chromatograms are scaled as 0-100% with 100% representing the ion current value listed in the upper right hand corner of the chromatograph (i.e., 1.89e6 for IL3-5 acylsucroses, panel A).

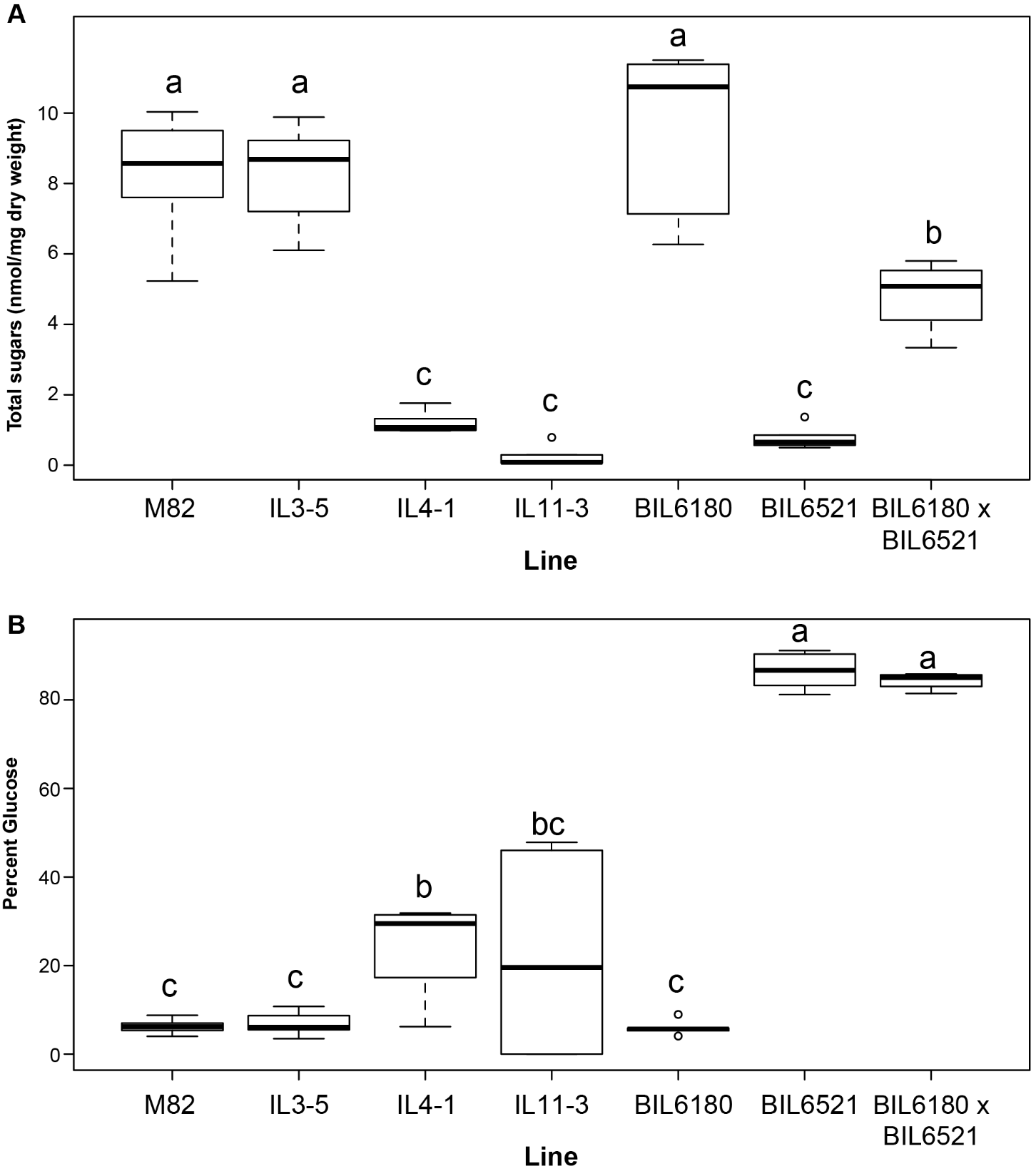

**Fig. S2.** **Quantification of acylsugars in *S. lycopersicum* M82 and breeding lines containing *S. pennellii* LA0716 introgressions.** (**A**) Total acylsugar accumulation quantified as the sum of sucrose and glucose from saponified acylsugar extracts. (**B**) Percentage of saponified sugars in acylsugar extracts detected as glucose. Treatments that do not share a letter are significantly different from one another (*p* < 0.05; one-way ANOVA, Tukey’s Honestly Significant Difference mean-separation test). Whiskers represent minimum and maximum values less than 1.5 times the interquartile range from the 1^st^ and 3^rd^ quartiles, respectively. Values outside this range are represented as circles; *n* = 6 for all ­lines except BIL6180 x BIL6521 F2 plants: *n* = 4.

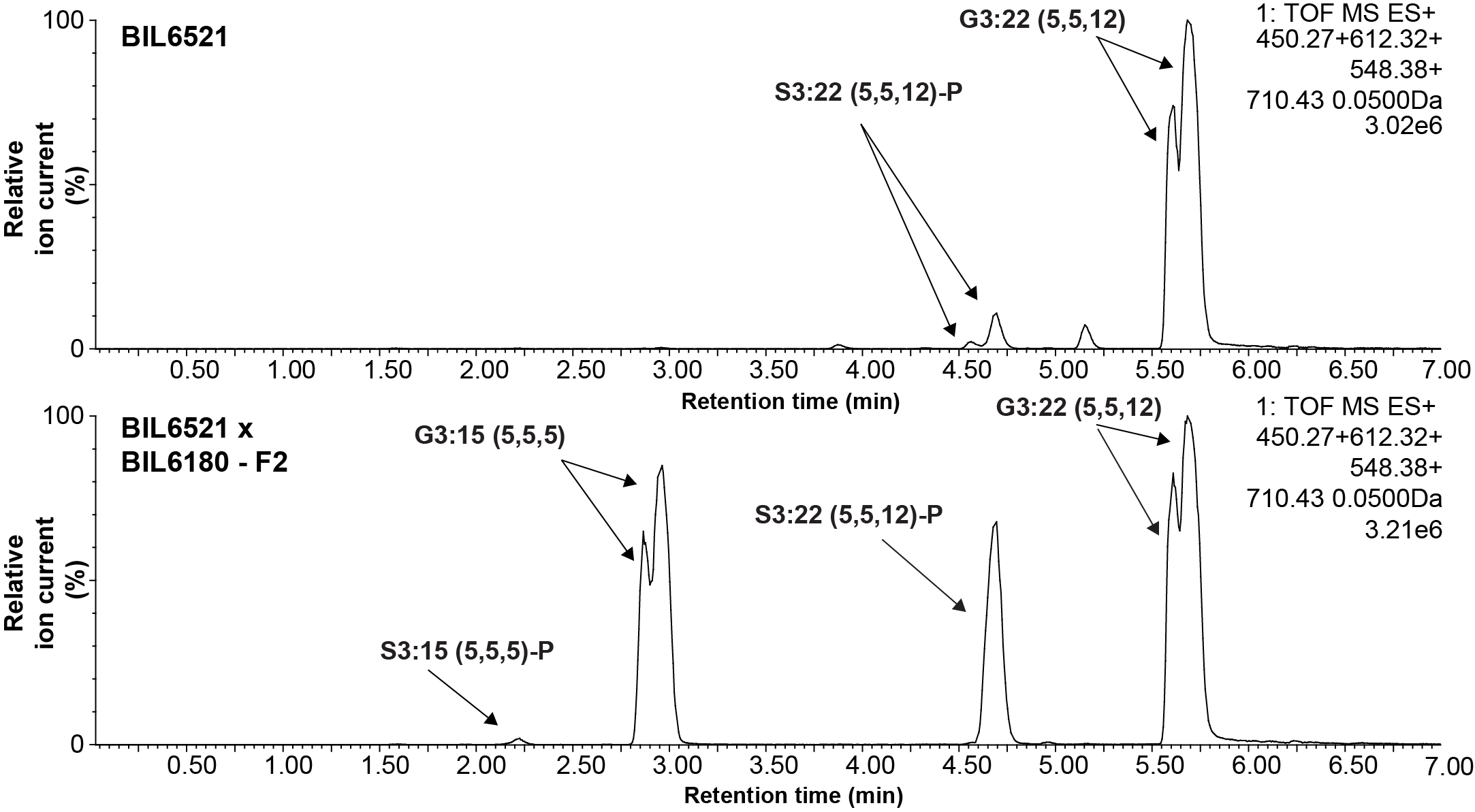

**Fig. S3. Comparison of major acylsugars in BIL6521 and BIL6521 x BIL6180 F2 progeny using LC-MS.** ESI+ mode LC-MS of trichome extracts reveals that the F2 progeny of BIL6521 crossed to BIL6180 (genotyped as heterozygous for Chr. 3 and 4 introgression regions) shows an increase in the short chain acylglucose, G3:15 compared to BIL6521 alone. Extracted ion chromatograms of NH_4_^+^ adducts of S3:15 (5,5,5) (*m/z*: 612.32) , G3:15 (5,5,5) (*m/z*: 450.27), S3:22 (5,5,12) (*m/z*: 710.43), and G3:22 (5,5,12) (*m/z*:548.38) in BIL6521 and BIL6521 x BIL6180 lines (mass window: 0.05 Da). Samples were run on the 7 min method described in the method section. The progeny were genotyped as described in the Methods section.

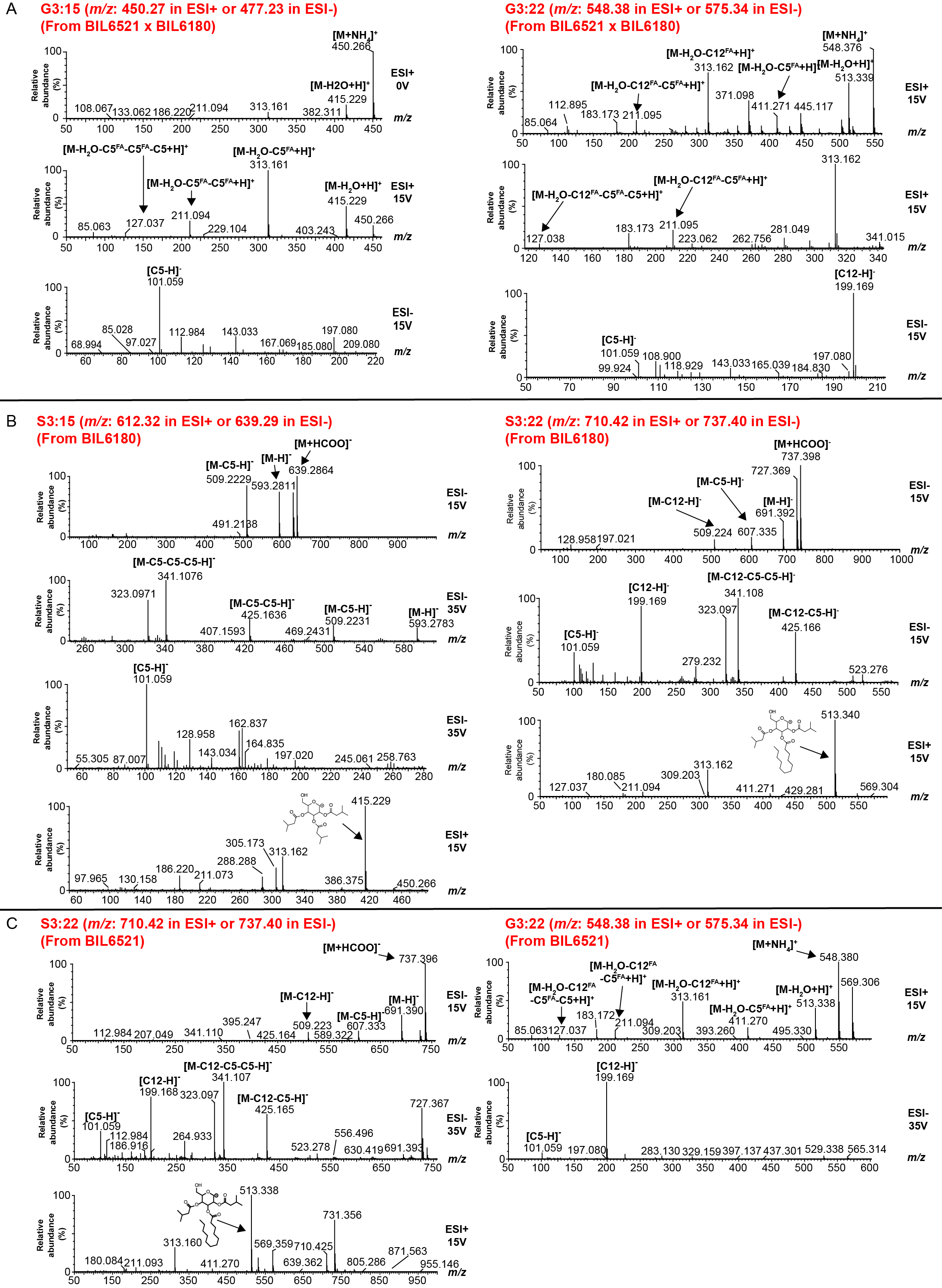

**Fig. S4. Mass spectra of major acylsugars S3:15, S3:22, G3:15, and G3:22 from BIL6521 x BIL6180 - F2 lines, BIL6180, and BIL6521.** Triacylglucoses fragment in either positive or negative ion mode by losing the first two acyl chains as neutral fatty acids followed by the third acyl chain lost as aliphatic ketene (R=C=O). In negative ion mode, fatty acid anions are also seen when the charge stays with the fatty acid fragment. Together these losses allow for determination of the length of acyl chains attached to the acylglucose. Acylsucroses fragment in negative ion mode with neutral losses of aliphatic ketenes. In positive ion mode, fragmentation of ammonium adducts of acylsucroses results in cleavage of the glycosidic linkage with the most stable (and most abundant) ion fragment coming from the charge staying on the furanose ring fragment. When no acyl chains are present on the furanose ring, the most abundant fragment ions are from the pyranose ring and further fragmentation results from neutral loss of fatty acids.(**A**) Fragmentation of G3:15 and G3:22 from BIL6521 x BIL6180. Fragmentation of G3:15 in ESI+ mode (0 and 15V) results in the loss of two C5 fatty acids, followed by loss of a C5 ketene. Fragmentation of G3:22 in ESI+ mode (15V) results in the loss of C12 and C5 fatty acid followed by loss of a C5 ketene. The higher collision energy at 15V ESI- reveals the presence of a C5 (*m/z:* 101.06*)*, or C5 (*m/z:* 101.06) and C12 (*m/z*: 199.17) fatty acids for G3:15 and G3:22 respectively. (**B**) Fragmentation of S3:15 and S3:22 from BIL6180. Fragmentation of S3:15 in ESI- mode (15 and 35V) is characterized by the loss of 3 C5 ketenes. Fragmentation of S3:22 in ESI- mode (15V) is characterized by the loss of one C12 ketene, and two C5 ketenes. C5 (*m/z*: 101.06) or C5 (*m/z*: 101.06) and C12 (*m/z*: 199.17) fatty acids are present in ESI- mode for S3:15 and S3:22 respectively. ESI+ mode (15V) of these two acylsugars reveals the presence of three C5 chains or two C5 chains and one C12 on the pyranose ring of sucrose of S3:15 and S3:22 respectively. (**C**) Fragmentation of S3:22 and G3:22 from BIL6521. The fragmentation of S3:22 in ESI- mode (15 and 35V) is characterized by the loss of one C12 and two C5 ketenes. ESI- mode (35V) also reveals the presence of C5 (*m/z*: 101.06) and C12 (*m/z*: 199.17) fatty acids. ESI+ mode (15V) fragmentation reveals all three acyl chains are present on the pyranose ring. Fragmentation of G3:22 in ESI+ mode (15V) is characterized by the loss of a C12 and C5 fatty acid, followed by the loss of a C5 ketene. ESI- mode (35V) reveals the presence of C5 (*m/z*: 101.06) and C12 (*m/z*: 199.17) fatty acids. All fragmentation in this figure was obtained using collision-induced dissociation of the acylsugars.

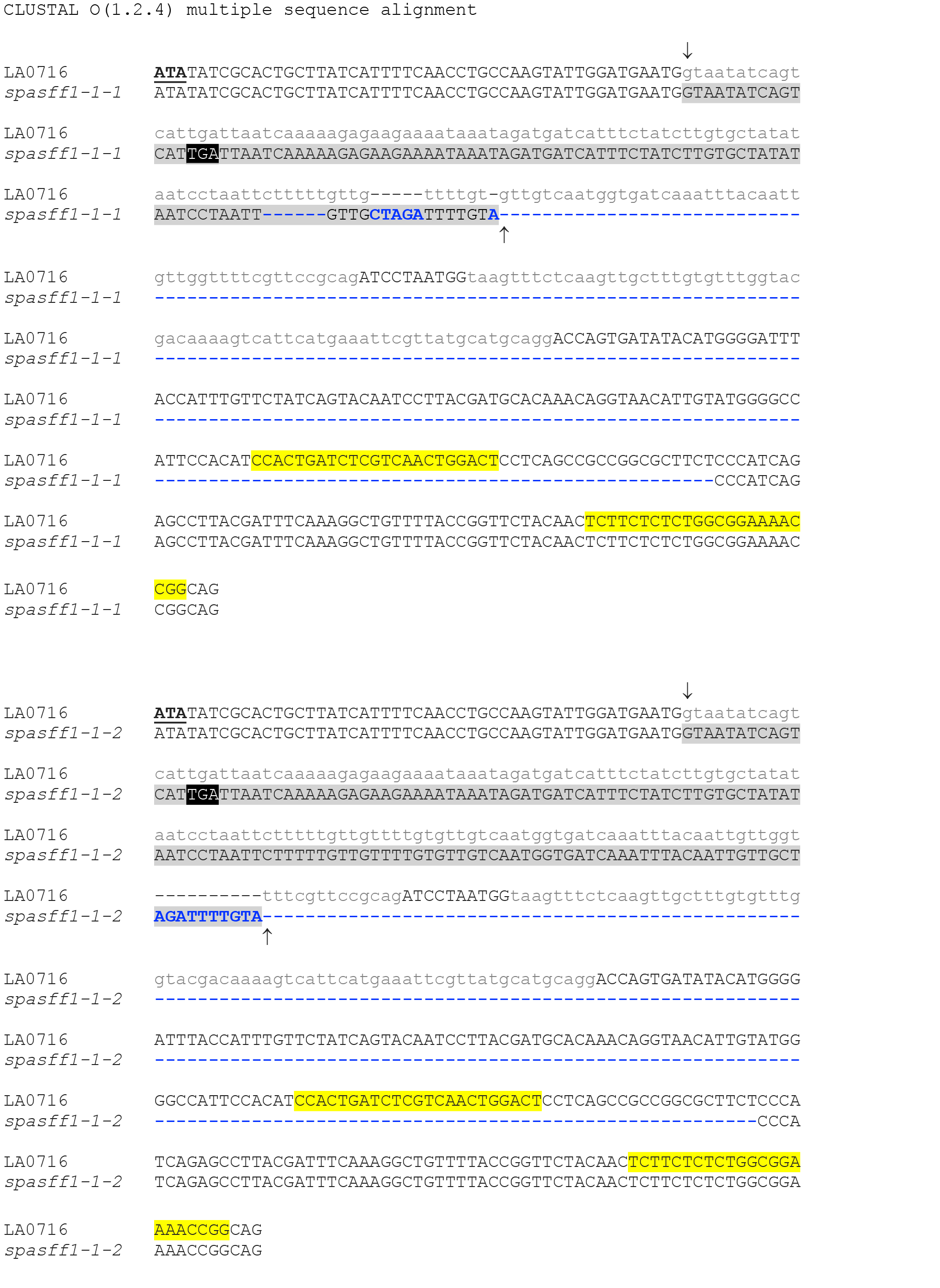

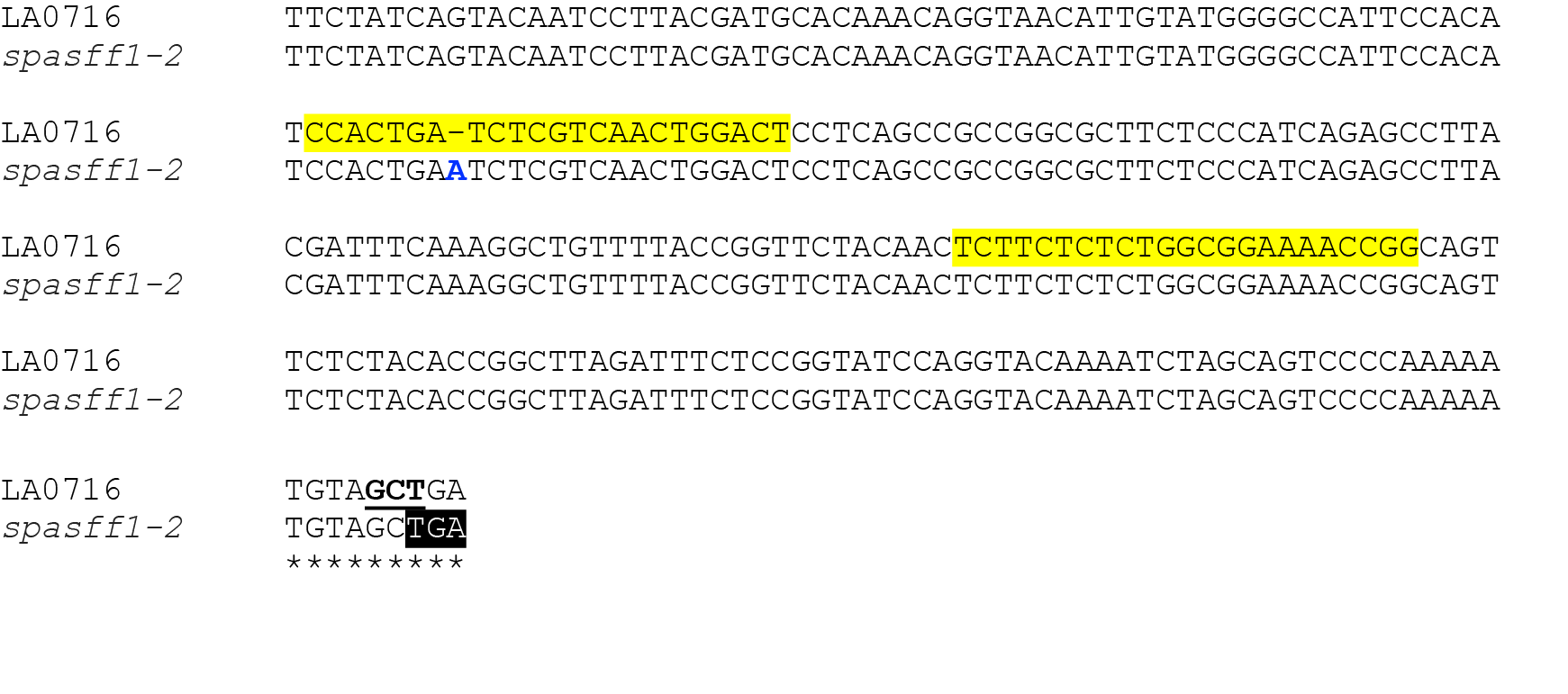

**Fig. S5. Mutated genomic sequence of three homozygous *spasff1* CRISPR/Cas9 lines.** Large insertion-deletions on *spasff1-1-1* and *spasff1-1-2* are shown by blue dashes and letters. Both mutations expand between exons (sequences shown in upper case and introns in lower case). We observed incorrect splicing events in transcripts in both mutant lines: arrows indicate splicing positions and extended exons in mutant lines are highlighted in grey. The mis-splicing events are predicted to result in premature stop codons, which are highlighted in black. Mutant *spasff1-*2 contains a single base pair insertion (blue letter) at one of the CRISPR/Cas9 target region (both regions are highlighted in yellow). This is predicted to cause a frame shift with resultant premature stop codon 180 nucleotides downstream (highlighted in black). The LA0716 wild-type allele reading frame is illustrated by a codon in bold and underlined.

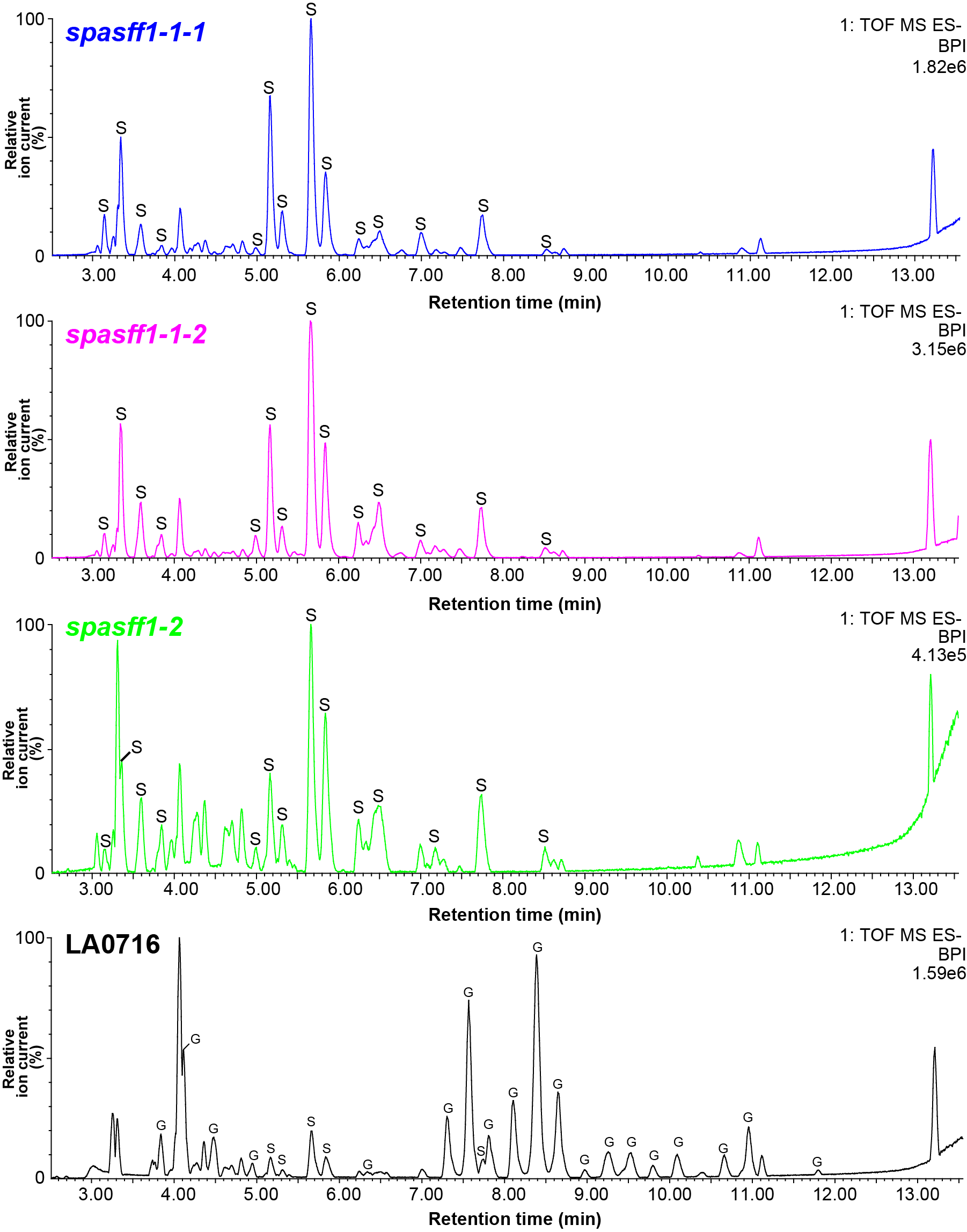

**Fig. S6. Acylsugars in BPI chromatograms of *spasff1* and LA0716 plants.** Base peak intensity (BPI) chromatograms of trichome extracts from *S. pennellii* LA0716 and the three *spasff1* lines shown in Fig 3C. The 21 min method and ESI- mode LC-MS was used for this analysis, as described in the Methods section. The 2.5 to 13.5 min is presented because acylsucroses and acylglucoses elute in this time period. Note: *spasff1* lines were diluted 100 fold before LC-MS analysis to avoid saturation of the LC-MS detector. This is due to differences in ionization between acylsucroses and acylglucoses in ESI- mode. Note: *spasff1-1-1/1-1-2* are homozygous T2 lines, while *spasff1-2* are homozygous T1 lines that were grown together (the same used in Fig. 3B and C).

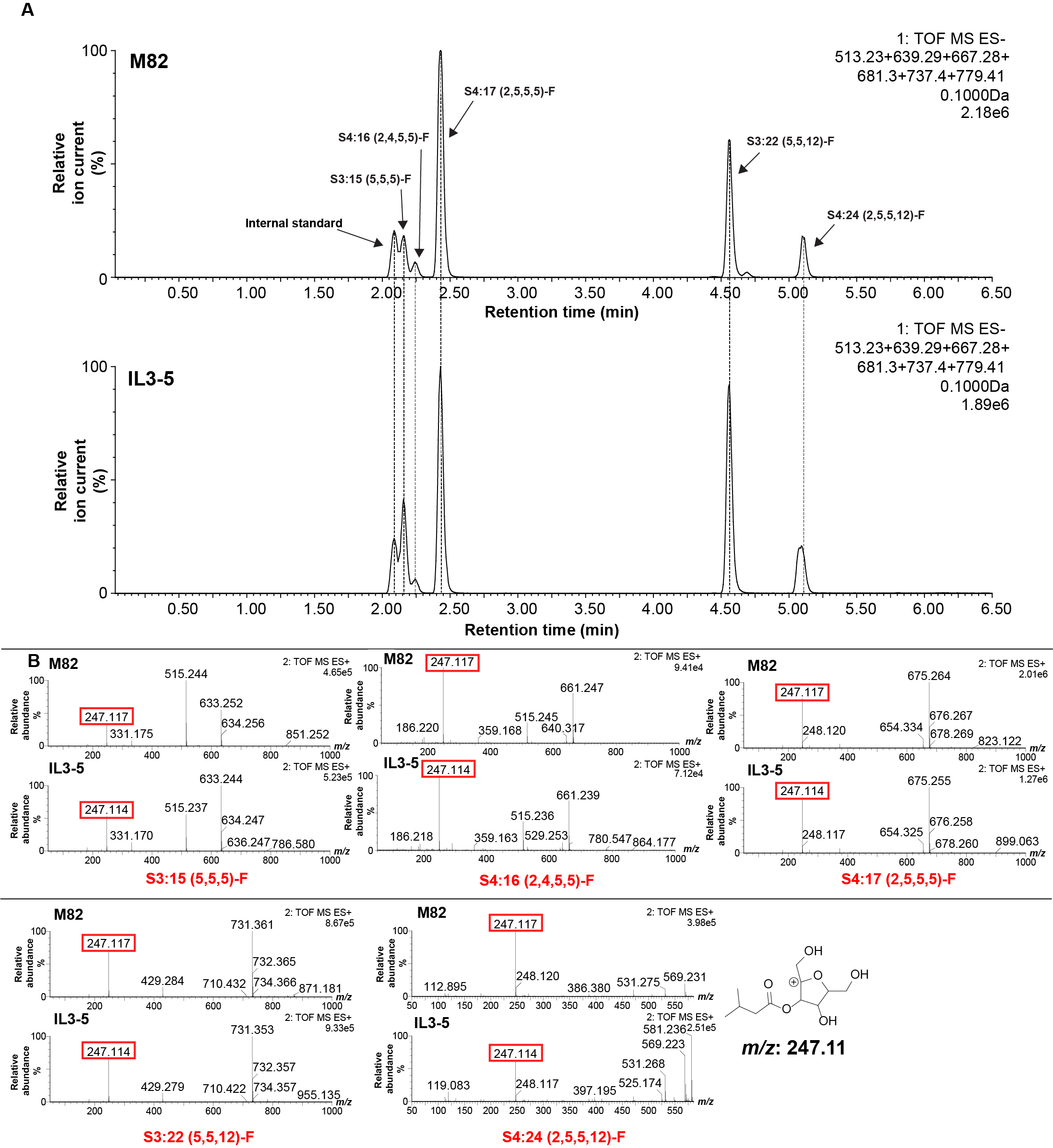

**Fig. S7. Comparison of acylsugars from IL3-5 and parental M82 using LC-MS.** (**A**) LC-MS analysis using ESI- mode to compare acylsugars of M82 and IL3-5. The major acylsugars in IL3-5 co-elute with acylsugars from M82. (**B**) ESI+ mode fragmentation of acylsucroses results in the cleavage of the glycosidic linkage (*8*). Fragment analysis of the acylsucroses using collision-induced dissociation from IL3-5 and M82 reveals a fragment ion (*m/z*: 247.11) consistent with the furanose ring of sucrose conjugated to a C5 acyl chain. This indicates that these acylsugars all possess a C5 acyl chain on the furanose ring. Samples were run on the 7 min method detailed in the Methods section (mass window: 0.1Da).

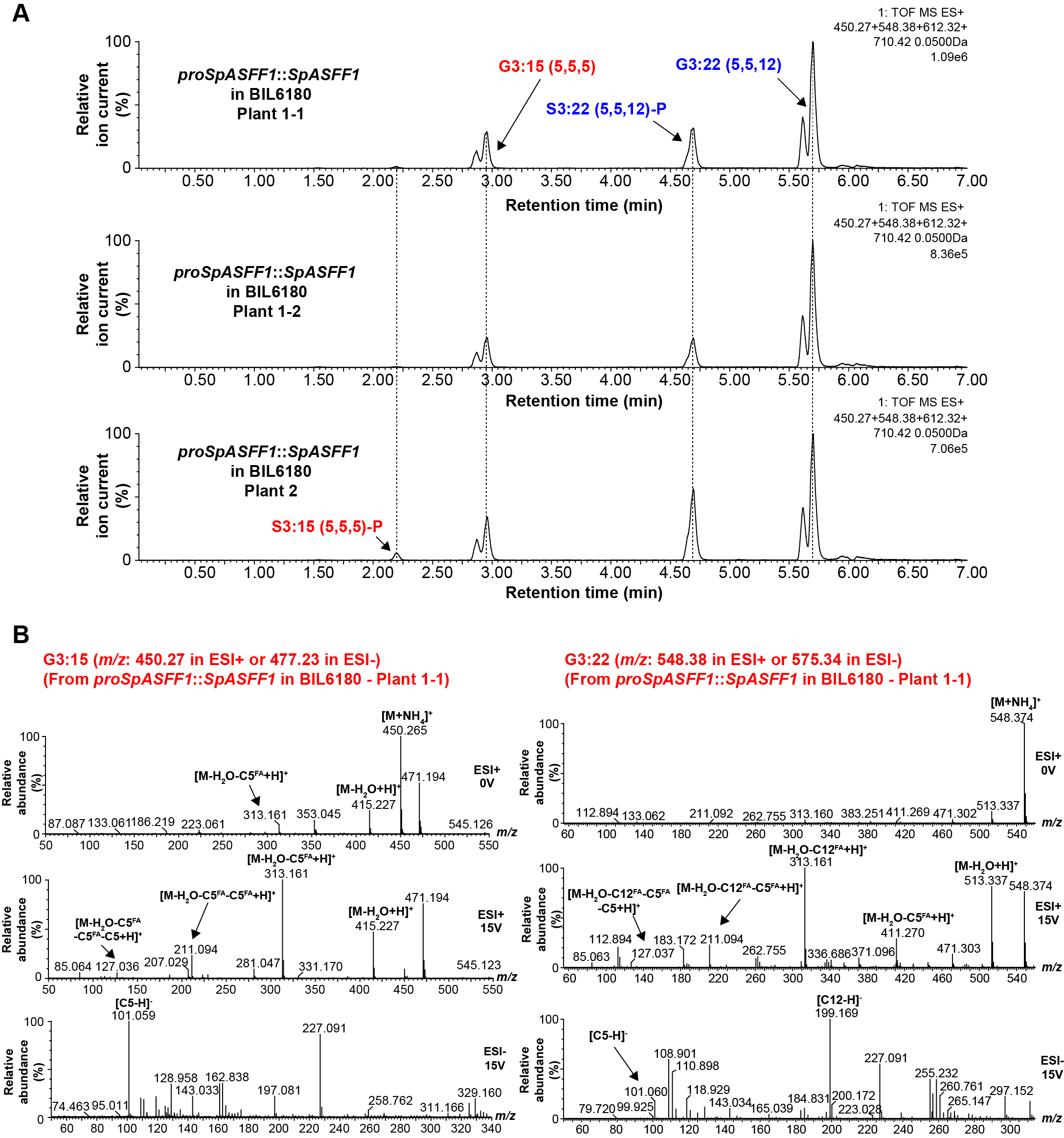

**Fig. S8. LC-MS analysis of P-type acylsucrose-producing *S. lycopersicum* BIL6180 stably transformed with *proSpASFF1*::*SpASFF1*.** (**A**) ESI+ mode LC-MS of trichome extracts from three *proSpASFF1*::*SpASFF1* T_2_ lines originating from 2 T_0_ lines. Extracted ion chromatograms of G3:15 (*m/z*:450.27), G3:22 (*m/z*:548.38), S3:15 (*m/z*: 612.32), and S3:22 (*m/z*: 710.42) are shown (mass window: 0.05 Da). Samples were run on the 7 min method described in the Methods section. Each pair of acylglucose peaks fragment similarly, consistent with the existence of alpha/beta acylglucose anomers. (**B**) Fragmentation of G3:15 and G3:22 from *proSpASFF1*::*SpASFF1* in BIL6180 – Plant 1-1 using collision-induced dissociation. Fragmentation of G3:15 in ESI+ mode (0 and 15V) results in the loss of two C5 fatty acids, followed by loss of a C5 ketene. Fragmentation of G3:22 in ESI+ mode (15V) results in the loss of C12 and C5 fatty acid followed by loss of a C5 ketene. The higher collision energy at 15V ESI- reveals the presence of a C5 (*m/z:* 101.06*)*, or C5 (*m/z:* 101.06) and C12 (*m/z*: 199.17) fatty acids for G3:15 and G3:22 respectively. Please reference Fig. S4 legend for further detail on fragmentation.

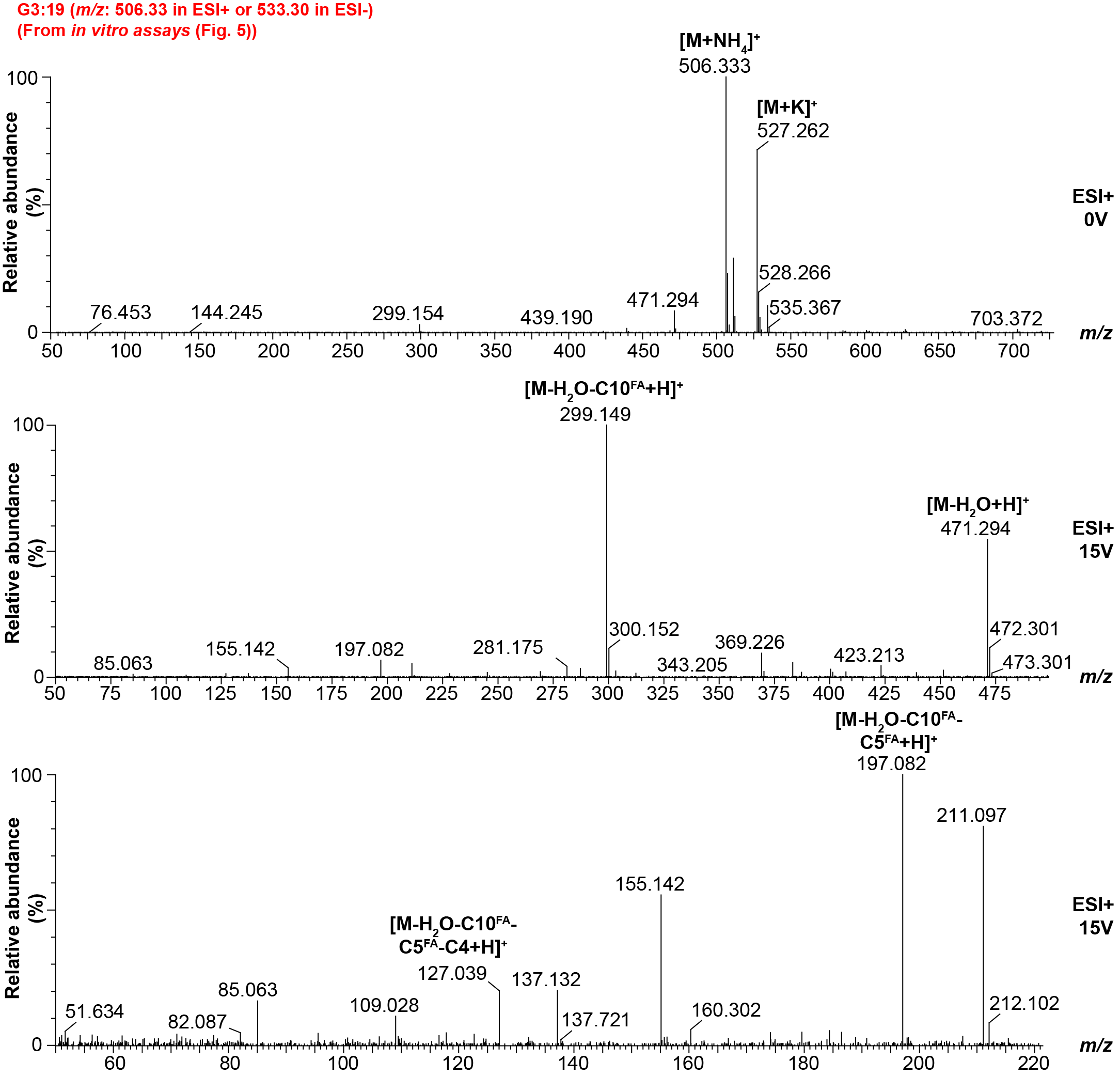

**Fig. S9.** **Mass spectra of G3:19-derived from SpASFF1 *in vitro* assay**. Fragmentation of the S3:19 + SpASFF1 reaction product, G3:19, in ESI+ mode (0V and 15V) using collision induced dissociation (from Fig. 5). The fragmentation is characterized by the loss of a C10 and C5 fatty acid from the triacylglucose, followed by loss of a C4 ketene. These results are consistent with the product cognate, S3:19. Further information of collision induced dissociation is present in Fig. S4.

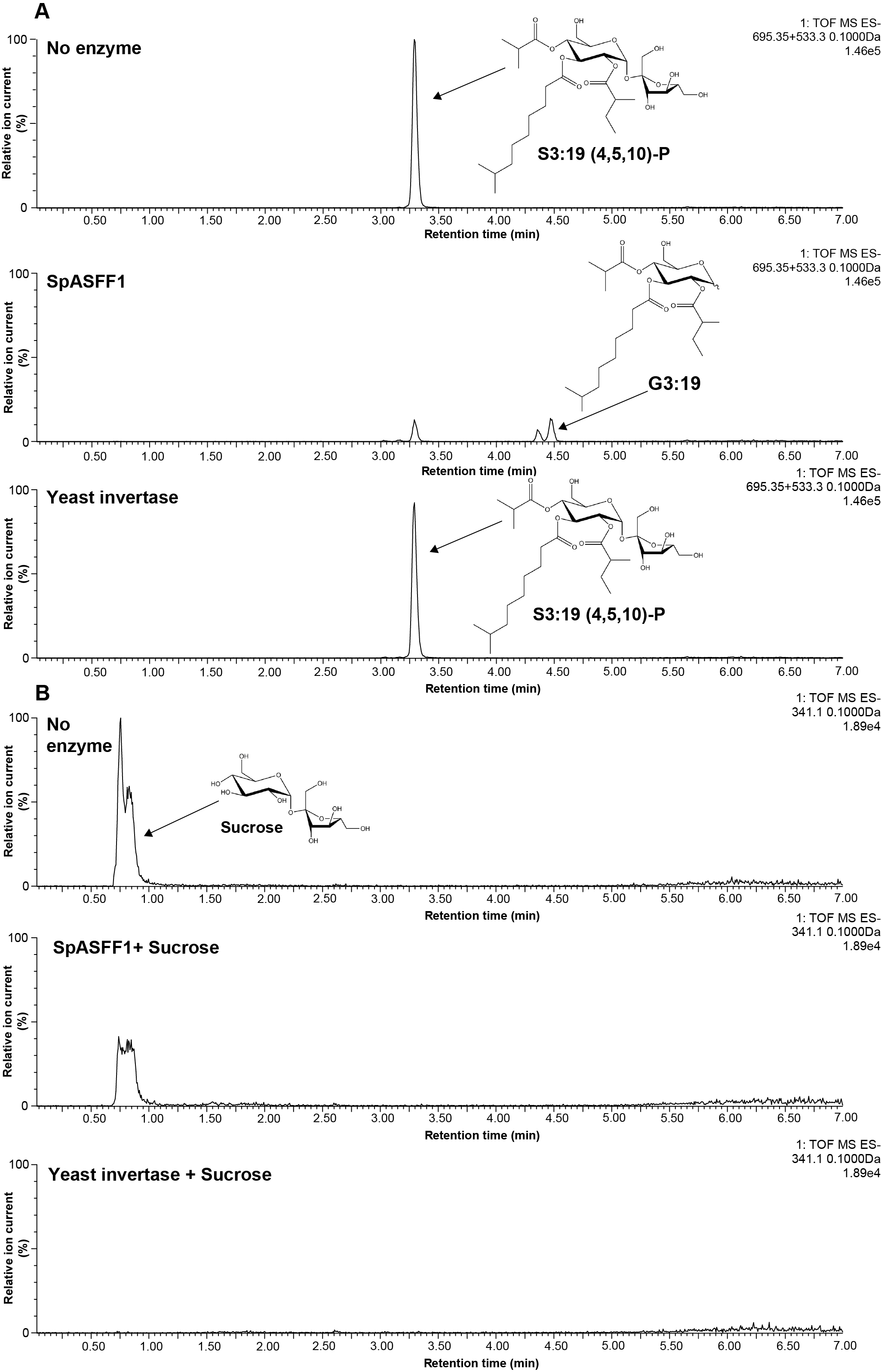

**Fig. S10.** **SpASFF1 cleaves a purified P-type triacylsucrose but not unmodified sucrose while yeast invertase cleaves unmodified sucrose but not triacylsucrose.** (**A**) LC-MS analysis of *in vitro* enzyme assay products indicates that SpASFF1 cleaves a P-type S3:19 (5^R2^,10^R3^,4^R4^) acylsucrose to yield a product with *m/z* = 533.3 (middle chromatograph) while yeast invertase yields no hydrolysis products (lower chromatograph). (**B**) LC-MS analysis of *in vitro* assays with unacylated sucrose indicates complete hydrolysis of sucrose by yeast invertase (lower chromatograph) but abundant sucrose remaining when incubated with SpASFF1 (middle chromatograph); disappearance of the sucrose substrate was monitored rather than appearance of the glucose or fructose products due to poor detection of the monosaccharides resulting from low ionization efficiency. Note: Acylglucose structure is inferred from collision induced dissociation-mediated fragmentation (Figure S9).

**Table S1. Annotation of acylsugars identified in BIL6521 and BIL6521 x BIL6180 F2 using LC-MS and collision-induced dissociation.** Listed acylsugars were identified from trichome extracts of BIL6521 and BIL6521 x BIL6180 F2 plants. Acylsugars were annotated using LC-qToF mass spectrometry in ESI-/+ mode using varying collision energies as described in Methods. Acylsugar fragmentation was analyzed as shown in Fig. S4 using collision-induced dissociation.

| BIL6521 acylsugars |  |  |  |  |
| --- | --- | --- | --- | --- |
| Mass/Charge ratio | ESI-/+ | Acylsugar annotation | Retention time (min) | Adduct |
| 681.30 | – | S4:17(2,5,5,5) | 2.44 | HCOO^-^ |
| 737.40 | – | S3:22 (5,5,12) | 4.56,4.69 | HCOO^-^ |
| 779.41 | – | S4:24 (2,5,5,12) | 5.10 | HCOO^-^ |
| 506.33 | + | G3:19 (5,5,9) | 4.84 | NH_4_^+^ |
| 520.34 | + | G3:20 (5,5,10) | 4.97 | NH_4_^+^ |
| 534.36 | + | G3:21 (5,5,11) | 5.23 | NH_4_^+^ |
| 548.38 | + | G3:22 (5,5,12) | 5.68 | NH_4_^+^ |
| BIL6521 x BIL6180 acylsugars |  |  |  |  |
| Mass/Charge ratio | ESI-/+ | Acylsugar annotation | Retention time (min) | Adduct |
| 639.29 | – | S3:15 (5,5,5) | 2.21 | HCOO^-^ |
| 681.30 | – | S4:17 (2,5,5,5) | 2.44 | HCOO^-^ |
| 723.38 | – | S3:21 (4,5,12) | 4.33 | HCOO^-^ |
| 737.40 | – | S3:22 (5,5,12) | 4.69 | HCOO^-^ |
| 751.41 | – | S3:23 (5,6,12) | 4.99 | HCOO^-^ |
| 436.25 | + | G3:14 (4,5,5) | 2.68 | NH_4_^+^ |
| 450.27 | + | G3:15(5,5,5) | 2.96 | NH_4_^+^ |
| 464.28 | + | G3:16 (5,5,6) | 3.25 | NH_4_^+^ |
| 520.34 | + | G3:20 (5,5,10) | 4.95 | NH_4_^+^ |
| 534.32 | + | G3:21 (5,5,11) | 4.56 | NH_4_^+^ |
| 548.38 | + | G3:22 (5,5,12) | 5.68 | NH_4_^+^ |

**Table S2. Annotation of three GH candidates for *SpASFF1* identified in the AG 3.2** This table provides the nominal or canonical activities of the three glycosyl hydrolase enzymes found in AG3.2 including members from the GH32, GH35, and GH47 families.

| **SGN ID**  **(*S. lycopersicum*/**  ***S. pennellii*)** | **Enzyme family** | **Nominal activity** | **Typical substrates** |
| --- | --- | --- | --- |
| Solyc03g121540/  Sopen03g040350 | GH35 | β-galactosidase | β-D-galactosides; α-L-arabinosides; oligo-galactosides (*47*) |
| Solyc03g121680/  Sopen03g040490 | GH32 | β-fructofuranosidase | β-fructofuranosides (*38*) |
| Solyc03g123900/  Sopen03g041640 | GH47 | α-mannosidase | Terminal α-D-mannose residues of oligo-mannose oligosaccharides (*48*) |

**Table S3. Primers/gBlocks/sgRNAs used in this study.** Red color indicates the NGG site in the sgRNAs.

| **Sequence name** | **Nucleotide sequence** |
| --- | --- |
| ASFF_F | ATGGGATATGTTAGAAGTGTTTGG |
| ASFF_R | TCAATTGATTTGAGCTGTTTTCA |
| (pEAQ-HT)-ASFF-His_F | GTATATTCTGCCCAAATTCGATGGGATATGTTAGAAGTGT |
| (pEAQ-HT)-ASFF-His_R | TGATGGTGATGGTGATGCCCATTGATTTGAGCTGTTTTCA |
| RT_actin_F | GGTCGTACCACTGGTATTGT |
| RT_actin_R | AAACGAAGAATGGCATGTGG |
| RT_ASFF_F | CTACGCAGGCAGATGTAGAAA |
| RT_ASFF_R | ATCACTAGAAGGCAAGTGTAAGG |
| RT_EF-1a_F | TGCTGCTGTAACAAGATGGA |
| RT_EF-1a_R | AGGGGATTTTGTCAGGGTTG |
| RT_ubiquitin_F | TCGTAAGGAGTGCCCTAATGCTGA |
| RT_ubiquitin_R | CAATCGCCTCCAGCCTTGTTGTAA |
| ASFF_001_F | CAAAAAAGCAGGCTCCGCCTGATAGTTATGCCAATGTACCACA |
| ASFF_001_R | ACGATCCTCTTAGTTGGTCCAC |
| ASFF_002_F | GTGGACCAACTAAGAGGATCGT |
| ASFF_002_R | TGGCCCCATACAATGTTACCTG |
| ASFF_003_F | CAGGTAACATTGTATGGGGCCA |
| ASFF_003_R | TTTGGGAAGTTCTGGCTCGG |
| ASFF_004_F | CCGAGCCAGAACTTCCCAAA |
| ASFF_004_R | GGGTCGGCGCGCCCACCCTTAAAGTGTTCGACTGACCATTCT |
| ASFF_Promoter_F1 | CAAAAAAGCAGGCTCCGCTGCCAATGTACCACAATTAGTATT |
| ASFF_Promoter_R1 | AGAAAGCTGGGTCGGCTTTTAGTTGAAGATGGCAACTACATTTCA |
| ASFF_Chr3_Indel_002_F | CAAAAAGAAGAAAAGGAAAACAGACA |
| ASFF_Chr3_Indel_002_R | GTGGGACTAAAACTTTGTAGTTCC |
| 04g011460_Marker_Indel-F | TAACAAAGCTTATGCACTCTTAG |
| 04g011460_Marker_Indel-R | ATCTACTACCTTCATATGCACAT |
| ASFF1_transcript_amp_01F | TCATTTCCATTCATAGCTATGGCA |
| ASFF1_transcript_amp_01R | GTTGCACCATCCACTGCTAA |
| sgRNA 1 target sequence w/ PAM site | AGTCCAGTTGACGAGATCAG**TGG** |
| sgRNA 2 target sequence w/ PAM site | TCTTCTCTCTGGCGGAAAAC**CGG** |
| sgRNA1 gBLOCK | TTTTAATGTACTGGGGTGGATGCAGTGGGCCCCACTCTGTGAAGACAAACTAGAATTCGAGCTCGGAGGGAGTGATCAAAAGTCCCACATCGATCAGGTGATATATAGCAGCTTAGTTTATATAATGATAGAGTCGACATAGCGATTGAGTCCAGTTGACGAGATCAGGTTTTAGAGCTAGAAATAGCAAGTTAAAATAAGGCTAGTCCGTTATCAACTTGAAAAAGTGGCACCGAGTCGGTGCTTTTTTTCTAGACCCAGCTTTCTTGTACAAAGTTGGCATTACGCTCGCTTTACTTGTCTTCTGCACGAAGTGGTTTAAACTATCAGTGTTTGACAGGATATATT |
| sgRNA2 gBLOCK | TTTAATGTACTGGGGTGGATGCAGTGGGCCCCACTCTGTGAAGACAATTACGAATTCCCATGGGGAGGGAGTGATCAAAAGTCCCACATCGATCAGGTGATATATAGCAGCTTAGTTTATATAATGATAGAGTCGACATAGCGATTGTCTTCTCTCTGGCGGAAAACGTTTTAGAGCTAGAAATAGCAAGTTAAAATAAGGCTAGTCCGTTATCAACTTGAAAAAGTGGCACCGAGTCGGTGCTTTTTTTCTAGACCCAGCTTTCTTGTACAAAGTTGGCATTACGCTCGCTCGCTCAGATTGTCTTCTGCACGAAGTGGTTTAAACTATCAGTGTTTGACAGGATATATT |
| pAGM4723_SeqF1 | TTTCGCCACCTCTGACTTGAG |
| pAGM4723_SeqR1 | CAGCTTGGCATCAGACAAACC |
| pICSL11024_SeqF1 | CGGCGAACTAATAACGCTCA |
| pICH47742CAS9_SeqF2 | ACTTACGCTCATCTCTTCGAC |
| pICH47742_SeqF1 | CTGTTGAATTACGTTAAGCATG |
| pICH41780_SeqR1 | TCGGTCACATGTGCATCCTC |
| ASFF_SeqR | GTTGATTTCCAACTACCATTCTCC |
| pK7WG-Kan-F | TTGACTCTAGCTAGAGTCCGAA |
| pK7WG-Kan-R | ATTGAACAAGATGGATTGCACGCA |
